# Ecological selection of dispersal strategies in metacommunities: impact of landscape features and competitive dynamics

**DOI:** 10.1101/2023.09.08.556897

**Authors:** Gabriel Khattar, Paul Savary, Pedro R. Peres-Neto

## Abstract

Dispersal is simultaneously a cause and a consequence of metacommunity dynamics. While the influence of dispersal on metacommunities is subject of intense research, we still do not understand how species-species and species-environment relationships determine the success of different dispersal strategies in metacommunities. To address this, we employed simulation models considering species with distinct context-dependent dispersal strategies involved in the three stages of dispersal (departure, transience, and settlement). These species were allowed to reach coexistence at the metacommunity scale under various competitive hierarchies and different levels of spatial and temporal environmental variability. By assessing the dispersal strategies of species that persisted and dominated metacommunities, we could understand how metacommunity dynamics impose ecological selection on dispersal. Our simulation model reproduced empirical patterns in species dispersal across different scales, ranging from changes in the success of dispersal strategies caused by local intraspecific and interspecific competition, to observed shifts in dispersal strategies along broad-scale ecological gradients. Additionally, we derived new empirically testable predictions regarding how metacommunity dynamics select for different dispersal strategies. Collectively, our results foster a comprehensive understanding of the factors influencing the success and diversity of dispersal strategies in a large array of ecological contexts.

## Introduction

Dispersal is the ecological process where individuals depart from their natal patches, move across the landscape, and eventually establish themselves in breeding patches. This multi-stage process regulates the spatial and temporal dynamics of natural systems across all levels of ecological organization [1,2]. At the metacommunity level, dispersal governs species coexistence [3] and influences community invasion success rates [4]. As a result, dispersal impacts diversity patterns within (alpha-diversity), between (beta-diversity), and across local communities (gamma-diversity).

Though much research has examined dispersal’s influence on metacommunity dynamics and diversity (see [5] and references within), our understanding of its role as a consequence of these dynamics is limited. For instance, density-dependent biotic interactions can regulate the decision of organisms to disperse or remain in their natal patches [6–8]. Similarly, resource availability [6], spatiotemporal heterogeneity [9,10], and landscapes’ physical connectivity [11] can impose ecological and evolutionary constraints on dispersal. The complex interplay between metacommunity dynamics and dispersal remains underexplored, yet discerning the forces governing species dispersal is crucial to understand the potential impacts of global change on biodiversity [12].

To understand how and why metacommunity dynamics should influence dispersal patterns in metacommunities, we need first to acknowledge the multi-stage nature and context-dependence of dispersal strategies, which are intentionally simplified in the foundational framework of metacommunity theory [10,13]. Dispersal arises from balancing decisions involving the timing to leave natal patches (i.e., emigration propensity), travelling distances (i.e., traversal), and the selection of a suitable new patch to settlement (i.e., habitat selection). This sequence of decisions determine the three stages of dispersal events, namely departure, movement, and settlement (*sensu* [14]). Dispersal decisions are context-dependent (plastic), meaning that organisms adjust them based on information about the surrounding biotic (e.g., predation, kin competition, and intra- and interspecific-competition) and abiotic (e.g., resource availability, spatiotemporal environmental variation) conditions [15]. For instance, organisms are more propense to leave their natal patches when local performance (i.e., fitness) is reduced via strong competition (intraspecific or interspecific), predation, resource scarcity, or unsuitable abiotic conditions [7,16]. These context-dependent decisions underly changes in species’ dispersal strategies to maximize regional fitness and/or minimize local mortality across different ecological contexts. Lastly, context-dependent variation in dispersal strategies are species-specific [6,8,16]. Thus, even species that are evolutionary closely related may still exhibit contrasting changes in dispersal strategies when subjected to varying abiotic and biotic conditions [8,16].

Given that metacommunity dynamics result directly from species-species and species-environment interactions, they should favor species from the regional pool whose dispersal strategies maximizes their persistence and dominance in the landscape [10]. For instance, species exhibiting a dispersal strategy characterized by a rapid increase in emigration propensity when local performance is decreased should have an advantage in persisting and dominating metacommunities in landscapes that undergo temporally variable habitat conditions [9]. Species that adopt a “risk-spreading” strategy, in which individuals choose to colonize suboptimal patches that have the potential to become optimal in the (relatively) short term, are expected to be favoured by temporal environmental variability. In metacommunities where competition dynamics hinder local coexistence (i.e., heterospecific competition is stronger than intraspecific competition [17]), species capable of reaching suitable habitat patches ahead of competitors have the potential to establish regional dominance through residency effects (*sensu* [18]).

Testing these (and potentially other) predictions about the ecological selective pressures of metacommunity dynamics on context-dependent dispersal strategies remains challenging for multiple reasons. Broad-scale observational data on multi-species dispersal strategies are scarce and experimental studies are commonly constrained by the number of species and environmental predictors that can be manipulated (but see [8,16,19]). Moreover, these studies are often conducted along a narrow range of ecological conditions, restricting their ability to capture the large range of conditions that could potentially drive variation in the success of distinct (context-dependent) dispersal strategies.

Here we sought out to expand our understanding about how landscape features and competition (ecologically) select for distinct context-dependent dispersal strategies in metacommunities (Figure 1). We employed process-based metacommunity models to assess how the interactions between landscape features and competition (i.e., metacommunity dynamics) select for dispersal strategies involving emigration propensity, habitat selection, and traversal (i.e., travelling distance). We allowed species with distinct context-dependent dispersal strategies to reach coexistence at the metacommunity scale under different types of competition types and under varying levels of spatiotemporal environmental variability (Fig. 1). By assessing the dispersal strategies of the species that persisted and dominated metacommunities, we were able to derive informed predictions regarding how metacommunity dynamics impose ecological selection on dispersal. Moreover, we demonstrated how the integration of species-specific context-dependent dispersal strategies into the basis of metacommunity theory can help us to understand the interdependence between community assembly and dispersal.

**Figure 1:**
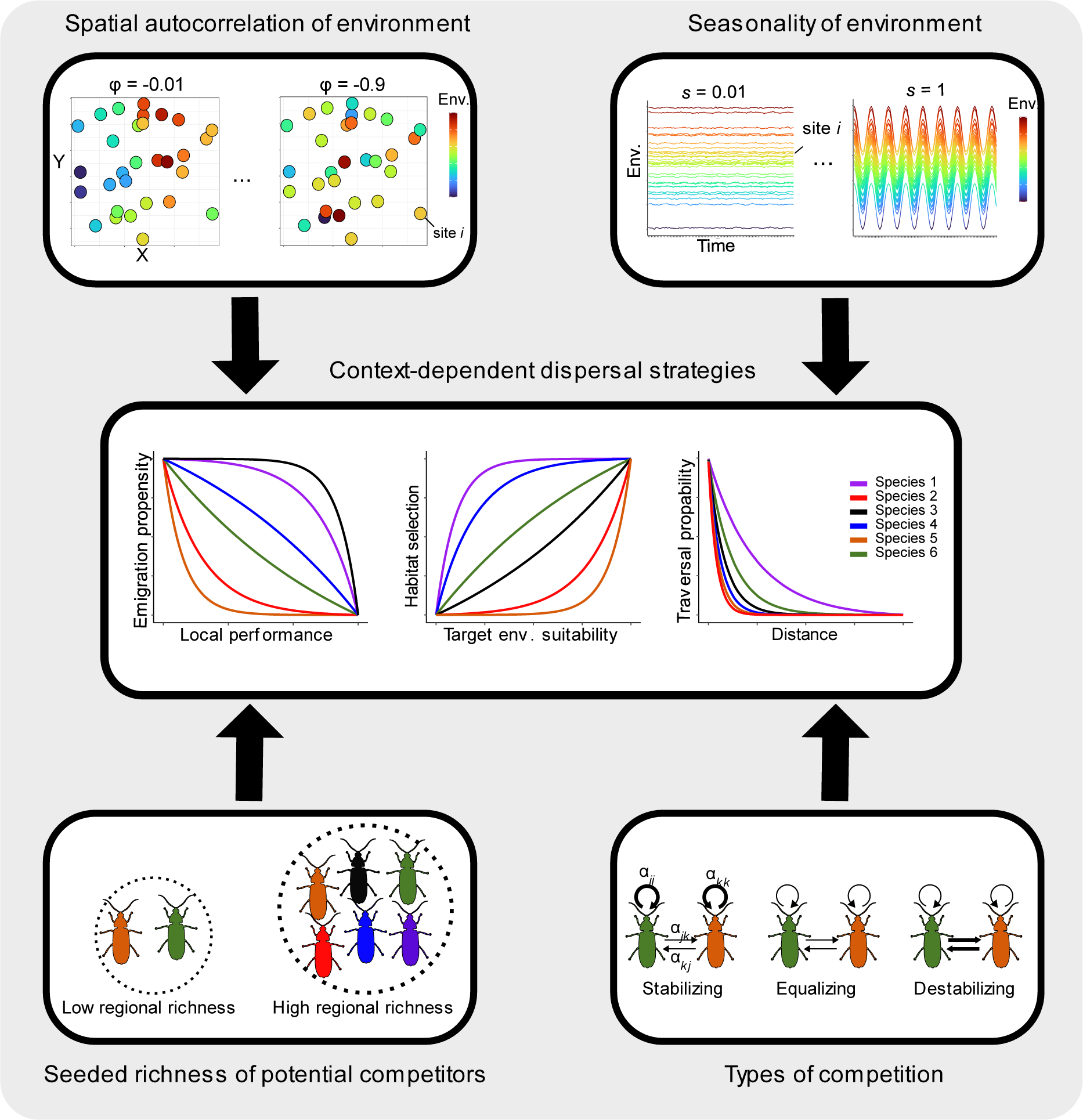
Simulation framework designed to understand how metacommunity dynamics impose ecological selection on context dependent dispersal strategies. Species were generated assuming random combinations of distinct dispersal strategies (i.e., represented by the plotted curves) for emigration propensity, habitat selection, and traversal probability (central panel). Species were allowed to colonise and reach coexistence in metacommunities subjected to different competitive dynamics (given by the factors represented in the lower panels) that took place in landscapes with different features (given by the factors represented in the upper panels). Parameters φ and s determine the landscape’s spatial structure in environmental conditions and seasonality, respectively (See more in Supp. Information I). The size of regional pools gives the richness of potential competitors at the beginning of each simulation iteration. Variation in competition type was simulated by manipulating the per-capita effects of a species on itself (intraspecific competition αkk, αjj) and on other species (interspecific competition αkj, αjk). Here we considered species pools under stabilizing (αkk = αjj > αkj= αjk), equalizing (αkk = αjj = αkj= αjk), and destabilizing (αkk = αjj < αkj= αjk) competition

## Methods

For the sake of brevity, we only briefly describe how we simulated landscapes and within-patch metacommunity dynamics here. An extended description and R code to recreate our models and further analyses are found in Supp. Material I and II.

### Simulated landscapes

We generated 25 landscape types following a cross-factorial design combining 5 levels of spatial autocorrelation (top-left panel in Figure 1) 5 levels of seasonality (top-right panel in Figure 1) in environmental conditions. We chose these two landscape features because they have been shown to impose costs and risks to species movement dictating ecological and evolutionary constraints on dispersal [9,10]. Each landscape type was composed of 50 habitat-patches with randomly generated x and y spatial coordinates (see Supp. Information I for more details).

### Parametrizing competitive dynamics in metacommunities

We seeded landscapes with species pools of different sizes (richness; 2 levels: 50 and 300 species) and assumed distinct types of competition among species (bottom-left and bottom-right panels in Figure 1). Species pool sizes affect metacommunity competition strength, as larger pools intensify competition for limited patches. We considered different types of competition by manipulating the per-capita effects of species on themselves (α_*intra*_) and their per-capita effect on other species (α_*inter*_) [13,20]. Stabilizing competition (α_*intra*_ > α_*inter*_) promotes local stable coexistence, enabling rare species to growth positively when populations of dominant competitors are in equilibrium. Under equalizing competitive dynamics (α_*intra*_ = α_*inter*_), individuals are neutral with respect to their per-capita effects on each other. This implies that coexistence is only possible for species having different niche requirements and/or display contrasting dispersal strategies. Under destabilizing competition (α_*intra*_ < α_*inter*_), local coexistence is unlikely because dominant species will lead rare species to local extinction even if those are better adapted to patch conditions.

#### Metacommunity dynamics

Our model simulates metacommunity dynamics that are spatially explicit, discrete in time, and governed by within-patch selection (density-dependent competition at the intraspecific and interspecific levels and density-independent species-environment sorting), dispersal, and ecological drift (as in [13]). Within-patch selection was modelled as a Beverton-Holt growth model [21] with generalized Lotka-Volterra competition assuming distinct competitive structures (i.e., stabilizing, equalizing, or destabilizing). An extended description of the model is found in Supp. Information I (see eq. SI-1 and SI-2).

Individuals able to persist in any given local community after within-patch selection and drift at time *t* could then disperse. Context-dependence in dispersal strategies was introduced by making species emigration propensity (*EP*), traversal probability (*TP*), and habitat selection (*HS*) to change as a function of local performance (given by joint influence of competition and niche-habitat matching), geographic distance, and environmental suitability, respectively. The shape of these relationships was made species-specific by randomly assigning different values of parameters *ep*, *tp*, and *hs* to each species in the regional pool.

Emigration propensity (*EP*) defines the probability that an individual leaves its natal patch given its current local performance. Based on previous experimental studies (e.g., [6]), we assumed that species were more propense to emigrate when local habitat conditions and/or biotic interactions decreased their local performance (*P*_*i,j,t*_, given by *eq SI-2*). As such, the emigration propensity of species *i* in site *j* at time *t* (*EP*_*i,j,t*_) decreased with local performance as follows:

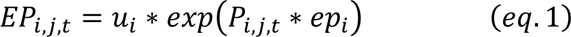

where *ep*_*i*_ is species-specific (equally spaced values within the interval [-0.1,0) U (0,0.1] that were then randomly assigned to each species), and determined the rate of change in *EP*_*i,j,t*_ as a function of *P*_*i,j,t*_ (scaled to range between 0 and 100). *u* determines the concavity of the relationship and was defined as:

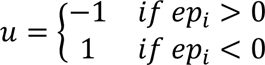

If *ep*_*i*_< 0 (and consequently *u* = 1), emigration propensity steeply decreased even at low levels of local performance (i.e., as observed in species 2, 5, and 6 in Figure 1, central panel). This strategy can be advantageous when local performance is temporally stable, allowing for competitive advantage due to residency effects. In contrast, species for which *ep*_*i*_> 0 (*u* = −1) were prone to emigrate from patches even if those provided high local performance (as observed in species 1, 3, and 4 Figure 1, central panel). This strategy can be advantageous when local performance changes abruptly due to environmental fluctuations. *EP*_*i,j,t*_(which is scaled to range between 0 to 1) set the probability of success in binomial trials determining the number of emigrants of species *i* departing from site *j* at time *t* (*E*_*i,j,t*_) after within-patch dynamics (given by the first term of *eq. SI-I*).

We relied on a random sampling process to determine the total number of immigrants of species *i* that will arrive at patch *j* at time t (*I*_*i,j,t*_) coming from other patches, e.g., patch *k* (*E*_*i,j,t*_). That is, let D = {*x, y*, …., *j*} be the potential destination patch for emigrants that departed from *k*. Let P = {*pi,kx,t, pi,ky,t,…., pi,kj,t*} be the set of probabilities for species *i* to immigrate to each patch in D when departing from *k* at time *t* (scaled to sum to unit). Note that *pi,kk,t* was set to 0 so that individuals that departed from *k* could not return to the same patch. Considering these two vectors, a random sampling process with unequal probabilities and replacement was repeated *E*_*i,j,t*_ times. The number of times patch *j* was sampled determined the number of individuals out of the total that departed from *k* (*E*_*i,k,t*_) that immigrated to patch *j* at time t (*I*_*i,kj,t*_). It follows that *I*_*i,j,t*_, i.e, the total number of individuals of species *i* that immigrated to *j* in time *t,* was then given by the sum of the total number of immigrants coming from all patches in the landscape.

We assumed that immigration probabilities increased with habitat suitability in the extant patch and decreased with the geographic distance between natal and extant patches. Thus, *pi,kj,t* was given by:

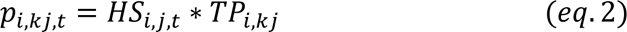

*HS*_*i,j,t*_ represents species *i* probability to move to patch *j* at time *t* based on the match between the environment in *j* (*Env*_*j,t*_) and their environmental requirements (i.e., *environmental suitability*, computed in the second term of *eq. SI. 2*). We assumed that species could assess the environmental suitability of extant patches at time *t* before deciding where to immigrate (i.e., informed dispersal). As such *HS*_*i,j,t*_increases with habitat suitability as follows:

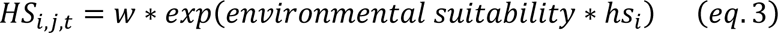

where *hs*_*i*_ is species-specific (equally spaced values within the interval [-0.1,0) U (0,0.1] that were then randomly assigned to each species) determining the rate of change in *HS*_*i,j,t*_ with *environmental suitability* (scaled to range between 0 and 100). *w* set the concavity and direction of this relationship and was defined as:

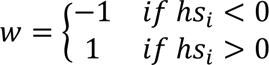

As such, species with *hs*_*i*_< 0 (and consequently a *w* = −1) displayed a risk-spreading strategy as they also tended to immigrate to patches with suboptimal habitat conditions (e.g., species 1, 4, and 6 in Figure 1, central panel). This strategy can be advantageous in landscapes with temporally variable habitat conditions. Conversely, species having *hs*_*i*_ > 0 (and consequently a *u* = 1) tended to immigrate only to patches with optimal or close to optimal environmental conditions (e.g., species 2, 3, and 5 in Figure 1, central panel). This strategy can be advantageous when environmental conditions are temporally constant across all patches.

*TPi*,kj is the probability of species *i* to move from patch *k* to *j* according to their geographic distance. *TPi*,kj decayed with the (Euclidean) distance between *k* and *j* (*dist*_*kj*_)as follows:

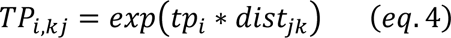

where *tp*_*i*_is species-specific (equally spaced values within the interval [-0.99, -0.01], that were then randomly assigned to each species) and determined the rate at which *TP*_*i,kj*_ decayed with *dist*_*kj*_. The smaller the *tp*_*i*_, the more limited the number of nearby patches a species could reach in a single dispersal event (e.g., species 2, 4, and 5, Figure 1 central panel). High traversal capacity (e.g., species 1, 3, and 6) should be favoured in landscapes where local habitat conditions are temporally variable, weakly autocorrelated in space, and when intraspecific competition is strong.

Lastly, we repeated the simulation framework described above considering two distinct assumptions about species ecological equivalence (see Supp. Information I for details). In one, species were set to exhibit distinct (non-neutral) performances along the environmental gradient. Then, species niche optima μ_*i*_took values equally spaced along the interval [0,5] to scale with environmental variation. Note that niche tolerance (σ) was set to be equal for all species and narrow enough to make them respond to environmental variation (see supplementary information). In the second assumption, we generated species to have equal (neutral) species’ responses (changes in performance) to environmental variation. This was operationalized by assigning the same environmental optima to all species (μ_*i*_ = average value of environmental conditions observed in the generated landscape). By contrasting simulation outputs between these two assumptions, we could investigate how species ecological equivalence can modulate (either buffer or amplify) the ecological selection of metacommunity dynamics on successful dispersal strategies.

#### Simulation iterations

Metacommunity dynamics were set to run for 1200-time steps (100 seasonal cycles). Each combination of scenarios was replicated 20 times, totalizing 6000 simulation iterations (20 replicates × 5 seasonality levels × 5 spatial structure levels × 2 species pool sizes × 3 competitive structures × 2 types of responses to environmental variation). At the first-time step (*t1*), we seeded each patch with species abundances randomly drawn from a Poisson distribution (λ = 0.5). Then, during the first ten seasonal cycles of each iteration (i.e., the first 120 time-steps), we seeded each patch with species abundances randomly drawn from a Poisson distribution with λ = 0.1. This seeding procedure ensured that species had equal chances to be initially present in all patches and that patches with similar environmental conditions could harbour different communities over time due to priority effects [13]. Metacommunity dynamics (with no seeding) ran for the remaining 1080 time-steps (i.e., 90 seasonal cycles) and we considered the metacommunity in the 100^th^ seasonal cycle (last 12 time-steps) for our analyses (see below). This ensured that model summaries were based on stable rather than transient metacommunities.

We estimated the most successful dispersal strategy as the metacommunity-weighted mean values of *ep*, *hs,* and *tp,* where the weights were given by a species’ regional abundance × occupancy (i.e., the relative number of patches occupied in the landscape). We also estimated the metacommunity-weighted standard deviation of *ep*, *hs,* and *tp* to quantify the “diversity” of dispersal strategies that resulted from metacommunity dynamics.

### Understanding how landscape features and competition dynamics select for dispersal strategies in metacommunities

We used random forest to assess how the metacommunity-weighted mean and standard deviation of *ep*, *hs*, and *tp* (continuous response variables) changed as a function of variation in seasonality (ordinal predictor, 5 levels), spatial autocorrelation (ordinal predictor, 5 levels), the seeded richness of competitors (ordinal predictor, 2 levels), different types of competition (categorical predictor, 3 levels), and different assumptions about species niche differentiation (categorical, 2 levels). Random forests identify general patterns in cross-factorial simulation data as they automatically model the effects of multilevel interactions among predictors on the response. We used the Boruta algorithm [22] to reduce model dimensionality and identify the most relevant predictors explaining variation in the metacommunity-weighted mean and standard deviation of *ep*, *hs*, and *tp*. Partial dependence plots revealed the direction of predictor-response relationships, controlling for other model predictors. We used bootstrapping to estimate 95% confidence intervals of predicted responses depicted in the partial dependence plots [23].

## Results and Discussion

Landscape features and competition dynamics had complex interactive effects on the success and diversity of dispersal strategies (i.e., metacommunity-weighted mean and standard deviation of *ep, hs, and tp*, respectively). Overall, the emigration propensity of the dominant species in the metacommunity decreased relatively fast with initial increases in local performance (i.e., *ep* < 0). Additionally, dominant species were highly selective for optimal conditions in destination patches (i.e., *hs* > 0) and could traverse large distances in a single dispersal event (i.e., ts > -0.5) (Fig. 2). Note though that optimal context-dependent dispersal strategies and their diversity changed consistently across levels of seasonality, environment’ spatial structure, competitors’ seeded richness, and competition types (Fig. 3 and Fig. SI. I). For the sake of tractability and synthesis, we focused on reporting and discussing the main effects of landscape features and competition types on the success and diversity of dispersal strategies. However, we also highlight and discuss some high-order interactions among predictors that increased our understanding of model outcomes (Figs SI. II-V).

**Figure 2:**
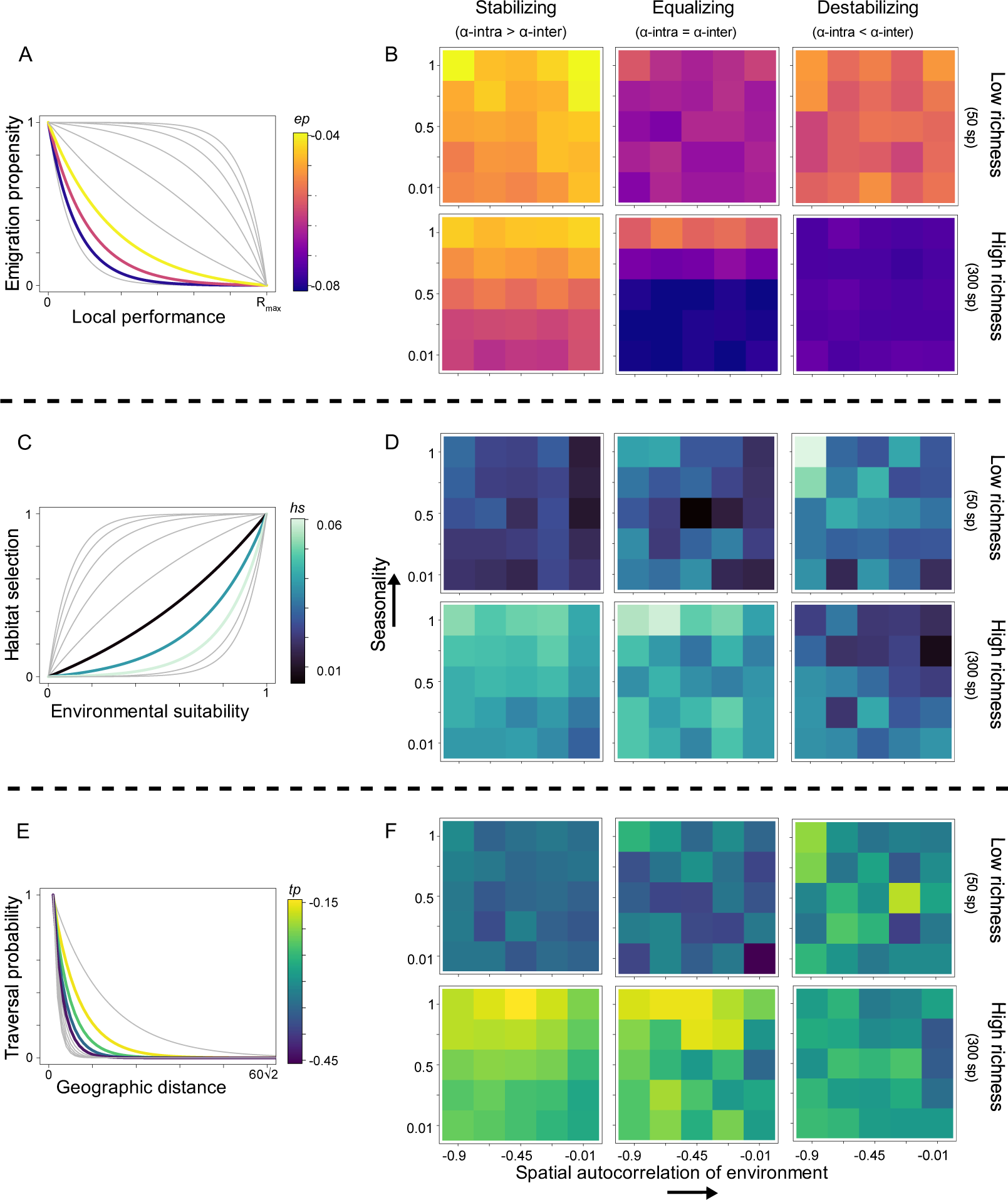
Dominant dispersal strategies for emigration propensity, habitat selection, and traversal probability observed across simulation scenarios. The curves in plots A, C, and E illustrate a small subset of the full range of context-depedent dispersal strategies seeded into metacommunities at each simulation iteration. The shape of these curves depends on the species-specific parameters ep, hs, and ts. The colour scales indicate the range of dispersal strategies that have dominated metacommunities at the end of each simulation iteration (estimated as the metacommunity-weighted mean values for hs, ep, ts). For illustrative purposes, some of these strategies are represented by the coloured curves in plots A, C, and E. The colour scales also serve as a reference for the heatmaps (B, D, and F) illustrating the changes in dominant context-dependent strategies across levels of seasonality, spatial autocorrelation, competitors’ richness, and competitive types. Each entry (square) in the heatmap represents the average value of the 20 metacommunity-weighted means obtained for a given simulation scenario. The results reported here were obtained under the assumption of niche differentiation (i.e., simulated species differed in habitat requirements). See Figs 3, SI-II, and S-IV for results obtained under the assumption of neutrality.

**Figure 3:**
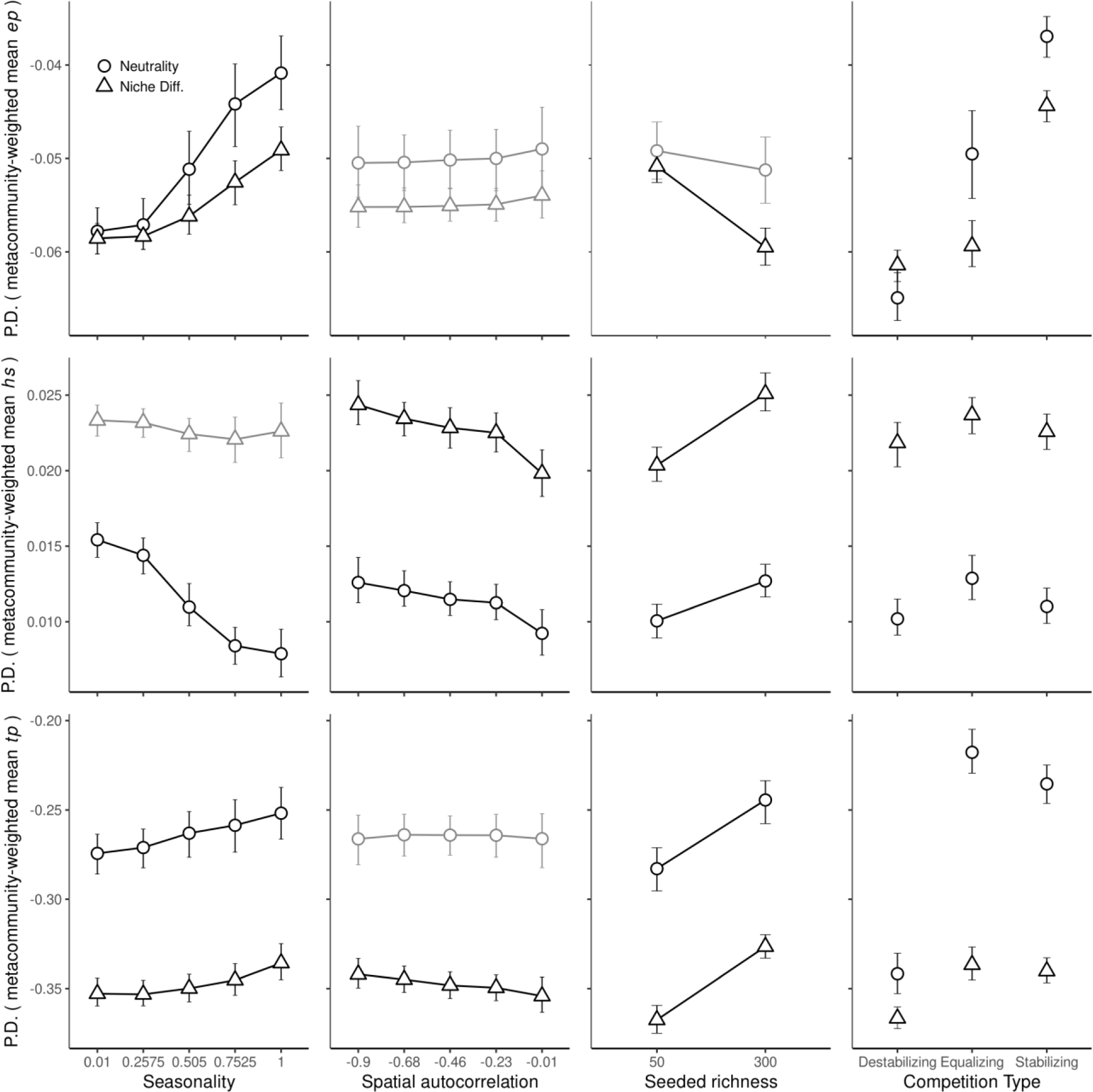
Partial dependence (PD) plots showing the predicted levels of dominant dispersal strategies in metacommunities (i.e., metacommunity-weighted mean values for ep, hs, ts) across levels of seasonality (first column), spatial autocorrelation in environmental conditions (second column), size of seeded regional pools (third column), and types of competition. Relationships were estimated for each level of the niche differentiation assumption (Neutrality = simulated species shared habitat requirements; Niche Diff. = simulated species differed in habitat requirements). Relationships of with variables that were not kept in the final random forests after feature selection were reported for illustrative purposes only (gray). R2 of random forest model fitted per row: metacommunity-weighted mean ep = 0.90, hs = 0.81, tp =0.95.

### The effects of landscape features on the success and diversity of dispersal strategies

In temporarily homogeneous (aseasonal) but spatially heterogeneous landscapes, species with lower average emigration rates (lower values of *ep*) were more successful in persisting and dominating metacommunities (Fig. 3 and Fig. SI. II). This because high emigration from temporally stable and high-performing patches can negatively affect species’ persistence, as reduced local abundances increases risks of local extinction from demographic stochasticity and competitive exclusion [24]. In contrast, when habitat conditions were seasonal, metacommunity dynamics favoured nomadic behaviours characterized by high emigration rates despite high levels of local performance. These “nomad” species were particularly favoured when they were also able to colonize several extant patches in single dispersal events (i.e., high *tp* values, Fig. 3 and Fig. SI. II). Essentially, species with high emigration rates and traversal capacity can rapidly shift their spatial distribution to cope with abrupt changes in local performance due to spatial and temporal variability in habitat conditions [9,25].

Empirical studies investigating latitudinal clines in species dispersal observed a similar influence of seasonality in selecting for higher emigration rates and traversal capacities. For instance, the relationship between seasonality and dispersal observed in our model serves as theoretical evidence that the conditions for the emergence of well-known latitudinal patterns on species range sizes and dispersal capacity (e.g.,[25,26]) can emerge by only considering metacommunity dynamics at fine spatiotemporal scales (i.e., no need to consider trait evolution and speciation, but see conclusions).

Note that when we assumed equivalent species’ environmental responses to environmental variation (i.e., neutrality), the relationship between seasonality and metacommunity-weighted mean *ep* and *tp* remained positive (Fig. 3 and Fig. SI. II). However, under this assumption, seasonality had a negative effect on the metacommunity-weighted mean *hs.* This implies that when species are neutral in relation to their habitat requirements, dispersing to temporally unsuitable patches, despite the risks, allows their persistence and dominance in seasonal landscapes. This is because colonizing briefly suboptimal patches enable them to broaden their spatial occupancy while avoiding intense competition.

In our model, the effect of spatial autocorrelation fostered insights into the influence of spatial uncertainty in habitat conditions on species’ dispersal strategies (Fig. 3 and Fig. SI. II). Species capable of minimizing dispersal risks by effectively tracking habitat conditions and reaching a larger number of patches (i.e., higher *hs* and *ts* values) were more successful in landscapes with weak spatial autocorrelation. In contrast, when habitat conditions were strongly autocorrelated in space, less selective species exhibiting weaker traversal capacity (i.e., lower *hs* and *ts* values) were more likely to persist in the metacommunity. Spatial autocorrelation modulates the respective success of dispersal strategies by determining the costs and risks of dispersal events. In landscapes with high spatial autocorrelation, the benefits of strong habitat selectivity and high traversal capacity diminish, as there are fewer unsuitable patches accessible from suitable ones.

Our results align with previous theoretical models, which observed that strong spatial autocorrelation favoured species with reduced traversal capacity, while weak autocorrelation favored species with strong traversal capacity [e.g., 13,39]. They also support empirical findings demonstrating that species adopt passive dispersal, a strategy characterized by weak habitat selection, when landscape structure is characterized by large clusters of patches with suitable habitat conditions [28].

It is important to note that when the niche differentiation assumption was relaxed, spatial autocorrelation had negligible effects on selecting for optimal traversal capacity strategy (Fig. 3 and Fig. SI. II). In our model, the neutrality assumption increases the number of species competing for a limited number of patches having equally suitable habitat conditions. As discussed below, a rise in species competing for similar habitat conditions favoured those with greater traversal capacity that enable them to escape local competition. Consequently, the selection for high-capacity traversers in highly competitive metacommunities offsets the selection of species with low traversal capacity in spatially structured landscapes (Figs. SI. II).

The diversity of dispersal strategies (weighted standard deviation of *ep*, *hs,* and *tp,* Fig. SI. I) also changed as a function of landscape seasonality and spatial autocorrelation (Figs. SI. I and III). Seasonality decreased the diversity of strategies related to habitat selection and traversal (i.e., the metacommunity-weighted standard deviation of *hs* and *tc,* respectively). Only highly selective species with long-distance traversal capacities could persist when local conditions fluctuated substantially over time. Alternatively, the diversity of emigration propensity strategies (i.e., metacommunity-weighted standard deviation of *ep*) increased with seasonality. Thus, although seasonality tended to favour species with relatively higher emigration rates, species with lower emigration rates could persist provided they accumulated enough individuals to buffer mortality in temporally unsuitable conditions (i.e., storage effects [17]).

Moreover, spatial autocorrelation in habitat conditions notably diversified traversal capacity and habitat selection strategies (Figs. SI. I and III). This is because large clusters of suitable habitat conditions decrease the risks of dispersing to unsuitable habitats which, in turn, facilitates the persistence of species with suboptimal strategies for traversal and habitat selection in the metacommunity.

### The effects of competition dynamics on the success and diversity of dispersal strategie

Research in metapopulation and movement ecology extensively explores how density-dependent processes [6,29], including intraspecific [30] and inter-specific competition [7] influence dispersal strategies. Consistent with these, our results indicate that increases in competition associated with larger species pools sizes favoured species with (i) reduced emigration propensity (lower *ep* values); (ii) pronounced selectivity towards habitat patch condition (i.e., higher *hs* values), and; (iii) strong traversal capacity (i.e., higher *hs* values) (Fig 3). These results held true across assumptions about niche differentiation and all types of competitive dynamics (Fig. SI.IV.)

Species with reduced emigration propensity were favoured at higher species richness levels because they ensured competitive dominance by maintaining large local populations (Figs 3 and SI.IV). The effectiveness of this strategy was maximized under destabilizing competition but minimized under stabilizing competition. Indeed, when intraspecific competition is stronger than interspecific competition, species with higher emigration propensity were able to mitigate the negative effects of intraspecific competition while keeping smaller viable populations in a larger number of suitable-habitat patches.

The success of dispersal strategies characterized by high traversal capacity increased with the initial number of competitors seeded in the metacommunity. This strategy had even greater success when intraspecific competition matched or surpassed interspecific competition (i.e., under equalizing and stabilizing competition, respectively, Fig SI. IV). Such trends resonate with empirical and theoretical studies that examine how population density, an indicative of intraspecific competition, impacts dispersal. Typically, these studies demonstrate that high densities increase emigration rates, particularly to patches farther away from their natal patch (see [31] and references within, and [30]). Our models shed deeper light on this dynamic, demonstrating that the relative strength of interspecific to intraspecific competition can either intensify or mitigate the latter’s impact on emigration rates and traversal.

We also observed that increasing the number of competitors favoured species that were more efficient in tracking and colonising suitable habitat conditions (higher *hs* values, Fig. 3). This efficiency was even greater under neutral competition dynamics (equalizing competition, Fig. SI.IV). Under equalizing competition, only species capable of tracking the limited number of patches where adequate niche-environment matching outweighs the negative effects of competitive interactions can coexist in the landscape. Taken together, species that were equal competitors but highly selective towards different habitats could coexist regionally through species-environment sorting dynamics.

Overall, the diversity of context-dependent dispersal strategies in the metacommunity decreased with the number of initial competitors seeded in the landscape (Figs. SI. I and SI. V). Thus, when a larger number of species competed for a few suitable patches, only a narrow range of dispersal strategies could ensure their regional persistence. Notably, the diversity of traversal capacity and habitat selection strategies (*tp,* and *hs*) did not follow this trend under destabilizing competition. In this case, only species with differences movement patterns in the landscape were able to coexist at the metacommunity scale by dominating distinct clusters of suitable patches [3]. In contrast, the diversity of dispersal strategies tended to increase when coexistence was facilitated through stabilizing competition.

### Conclusions, assumptions, and future directions

In this study, we used models to demonstrate that dispersal not only shapes the structure of metacommunities but also emerges from metacommunity dynamics. Previous theoretical and empirical studies that shared similar goals focused on investigating the ecological drivers of fixed behaviours involved in one or two stages of dispersal [10]. Our study stands out as the first to use metacommunity theory to generate predictions regarding the selective effects of landscape features and competition dynamics on species-specific context-dependent dispersal behaviours involved in all three dispersal stages (i.e., departure, transience, and settlement). Our models effectively recreated well-known variations in dispersal patterns across spatial scales, including changes caused by different forms of intraspecific and interspecific competition at local spatial scales and also shifts in dispersal patterns along broad-scale ecological gradients. Thus, our study improves the understanding of the factors influencing the success and diversity of dispersal strategies in a large array of ecological contexts.

However, our simulation models did not encompass the full complexity of species dispersal and its intricate relationships with metacommunity dynamics. For instance, we did not consider cost-related trade-offs that can cause covariance between dispersal, morphological, and behavioural traits. For instance, colonization-competition and ecological specialization-dispersal trade-offs can emerge as eco-evolutionary consequences of community assembly in landscapes with varying levels of environmental stability and habitat heterogeneity [17,32]. Therefore, we should expect these dispersal trade-offs to also be selected by the metacommunity dynamics.

Moreover, we focused solely on how competition at both intra and interspecific levels affects dispersal patterns in metacommunities. Yet, empirical experimental studies suggest that other biotic interactions can select for optimum context-dependent dispersal strategies. For example, predation risk can drive emigration [6],while parasitism can have dual effects on host dispersal: it can stimulate movement if the host perceives threat and relocates, or it can inhibit movement if the host stays and becomes infected [29].Incorporating these and other biotic interactions into our framework is a logical progression for future research.

Lastly, in our model, although we investigated how biotic and abiotic factors selected for a wide range of predefined traits (*ep, hs* and *tc*) that determine species-specific dispersal strategies, we did not consider trait evolution at the species level. Trait evolution is an important component of metacommunity theory [33,34] and the literature includes examples where the role of dispersal for maintaining biodiversity in changing environments is counteracted by the evolution of traits determining species’ habitat requirements [35]. Moreover, we also did not consider the full range of possible non-linear changes in dispersal strategies commonly observed in nature. For instance, we assumed that emigration propensity decreased monotonically and continuously with local performance. However, Allee effects or other density-dependent behaviours could lead to a wide range of non-linear relationships between population density, and hence local performance, and dispersal (e.g., u-shaped [7] or threshold functions [36]). To address these limitations, future studies could build upon our modelling framework to explore how trait evolution and a wider range of dispersal strategies are modulated by landscape features and competition dynamics.

## Supporting information

Code and Data to run simulations and generate figures

## Acknowledgements

We are grateful to Emanuel A. Fronhofer for insightful comments on the initial drafts of this manuscript. GK is supported by the Canada Research Chair in Spatial Ecology and Biogeography held by PPN. PS is funded by a Concordia Horizon postdoctoral fellowship and the Canada Research Chair in Spatial Ecology and Biodiversity held by PPN

## Notes

### Competing Interest Statement

The authors have declared no competing interest.

