## Supplemental Information for "Ecological selection of dispersal strategies in metacommunities: impact of landscape features and competitive dynamics"

**Supplementary Information**

**Simulated landscapes: Extended description**

We generated 25 landscape types following a cross-factorial design combining 5 levels of seasonality (top-right panel in Figure 1) and 5 levels of spatial autocorrelation (top-left panel in Figure 1) in environmental conditions. We chose these two landscape features because they have been shown to impose costs and risks to species movement that ultimately dictate ecological and evolutionary constraints on dispersal [1,2].

We started by assigning 50 patches into the landscape by drawing their x and y spatial coordinates from a uniform distribution ranging from 0 to 60. Environmental conditions at the patch level were defined by a single continuous variable ($Env$), spatially autocorrelated across patches and bound within the interval [0,5]. Spatial habitat autocorrelation in the environment was generated by random draws from a multivariate normal distribution (mu=0) with a covariance matrix set as the desired spatial autocorrelation level. Covariance matrices were created such that environmental similarity decayed exponentially with geographic distance according to *φ*. By manipulating the levels of *φ,* we generated landscapes where environmental conditions ranged from weakly autocorrelated (e.g., *φ* = -0.9) to strongly autocorrelated (e.g., *φ* = -0.01).

To simulate seasonal environmental variation, we set local environmental conditions to follow a sinusoid function with 100 periods, each composed of 12-time steps (i.e., 100 years) plus a random error $N\left( 0,0.1 \right)$ that served to mimic the effects of temporal environmental stochasticity. The amplitude of the sinusoidal environmental variation was modulated by a multiplicative factor *s* (constant across all patches). By manipulating the values of *s*, we created landscapes ranging from highly aseasonal (*s* = 0.1) to highly seasonal (*s* = 1).

We then seeded metacommunity simulations (see *Metacommunity dynamics* below) with the final matrices containing environmental values of each patch over time ($\mathbf{Env}$, 50 patches x 1200 times), and their corresponding spatial coordinates.

***Metacommunity dynamics: Extended description of within-patch mechanisms***

The dynamics of ecological communities were spatially explicit, discrete in time, and governed by habitat selection, demographic stochasticity, competition at intra and interspecific levels, and dispersal [3,4].

Considering that *N_i,j,t_* is the abundance of species *i* in site *j* at time *t*, population dynamics is governed by:

$N_{i,j,t}=Poisson\left( N_{i,j,t-1}*P_{i,j,t} \right)-E_{i,j,t}+I_{i,j,t}$ (eq. SI-1)

The first term of eq. 1 is a modified version of the Beverton-Holt competition model [5] that sets population growth as a function of habitat selection, competitive dynamics, and demographic stochasticity. $E_{i,j,t}$ and $I_{i,j,t}$ are the total number of individuals of *i* that emigrate from and immigrate to site *j* at time *t*, respectively.

$P_{i,j,t}$, is the local performance (i.e., growth rate) of species *i* when conditioned to competition and habitat selection and is modelled as follows:

$P_{i,j,t}=R.max*exp\left( \frac{-\left( {Env}_{j,t}-\mu_{i} \right)^{2}}{2\sigma^{2}} \right)*\frac{1}{\left( 1+\alpha_{intra}N_{i,j,t}+\alpha_{inter}\sum_{k\neq i}^{S} N_{k,j,t} \right)}$	(eq. SI-2)

$R.max$ represented the species' maximum intrinsic growth rate and was assumed to be equal to 3 for all species. Making $R.max$ equal across species ensured that they could reach the same maximum growth rate when optimum habitat conditions and competition at the intra and interspecific levels are negligible. $P_{i,j,t}$ ranges from 0 to $R.max$

The second term of eq. SI-2 sets *environmental suitability, i.e.,* the match between local environmental conditions ${( Env}_{jt})$ and species' environmental requirements. *environmental suitability* values ranges between 0 to 1, representing a complete mismatch and a complete match between species niche optima and local habitat conditions, respectively . We simulated species exhibiting distinct (non-neutral) performances along the same environmental gradient. To that end, species niche optima $\mu_{i}$ took values equally spaced along the interval [0,5] to scale with environmental variation. Note that niche tolerance $\left( \sigma\right)$ was fixed across all species and narrow enough to make species respond to environmental variation. Preliminary simulations showed that fixing $\sigma=1$ makes species sensitive to environmental variation without making them overly prone to local extinctions when conditions are suboptimal.

In parallel, to investigate how the assumption of niche differentiation can modulate (either buffer or potentialize) the influence of metacommunity dynamics on successful dispersal strategies, we also ran simulations with equal (neutral) species' responses to environmental variation. This was operationalized by assigning the same environmental optima to all species ($\mu_{i}$ = average value of $\mathbf{Env}$).

The third term of eq. SI-2 models the effects of density-dependent competition on population dynamics. Stabilizing competition was set as $\alpha_{intra}$= 0.0066 > $\alpha_{inter}=0.0033$; equalizing competition as $\alpha_{intra}$ = $\alpha_{inter}=0.005$; destabilizing competition as $\alpha_{intra}$ = 0.0033 < $\alpha_{inter}=0.0066$. These values imply the assumption that when locally dominant species make up 50% of the total community abundance, they face the same level of per-capita competition across the three types of competition. Prior tests (not shown) demonstrated that relaxing this assumption by considering different combinations of values for $\alpha_{intra}$ and $\alpha_{inter}$ did not influence the observed patterns of local coexistence and exclusion. However, they did alter the abundances at which locally coexisting species reached stable equilibria.

To incorporate the influence of demographic stochasticity on local birth and survival, we draw the final local species abundances from a Poisson distribution whose mean was determined by the estimated population size after habitat suitability and density-dependent competition [3,6].

Individuals able to persist in any given local community after within-patch selection and drift at time *t* could then disperse. See main text for a detailed description of how dispersal (departure, movement, and settlement) was computed.

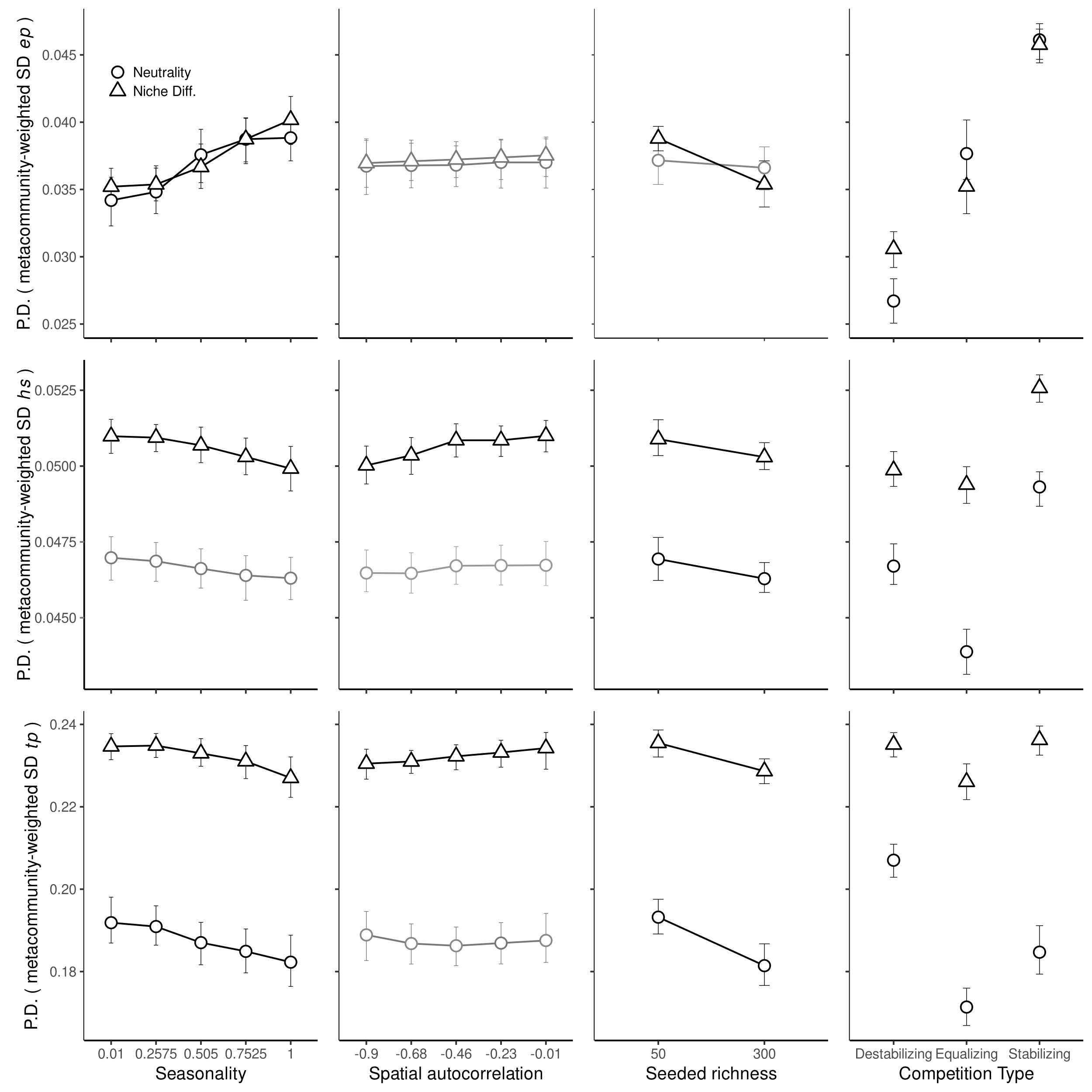

Figure SI. I: Partial dependence (PD) plots showing the predicted diversity of dispersal strategies (i.e., metacommunity-weighted standard-deviation for ep, hs, ts) across levels of seasonality (first column), spatial autocorrelation of environmental conditions (second column), size of seeded regional pools (third column), and types of competition. Relationships were estimated for each niche differentiation assumption (Neutrality = species shared habitat requirements; Niche Diff. = species’ habitat requirements differed). Relationships of ep, hs, and ts with variables that were not kept in the final random forests after feature selection were reported in gray for illustrative purposes only. R2 of random forest model fitted per row: metacommunity-weighted standard deviation ep = 0.89, hs = 0.75, tp =0.9.

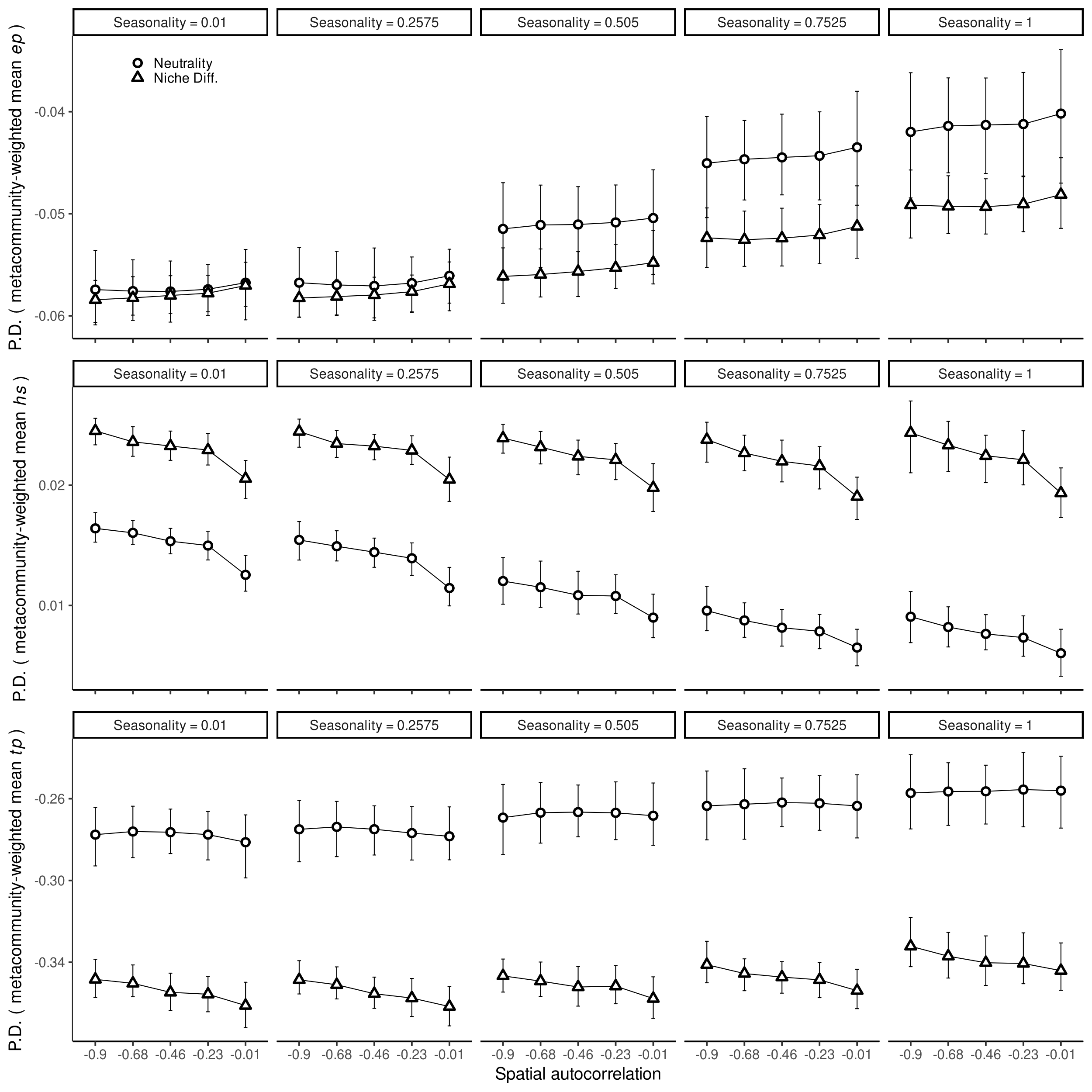

Figure SI. II: Partial dependence (PD) plots showing interactive effects of seasonality (panels), spatial autocorrelation (x-axis), and the niche differentiation assumption (symbols) on the dominant dispersal strategies in metacommunities (i.e., metacommunity-weighted mean values for *ep, hs, ts).*

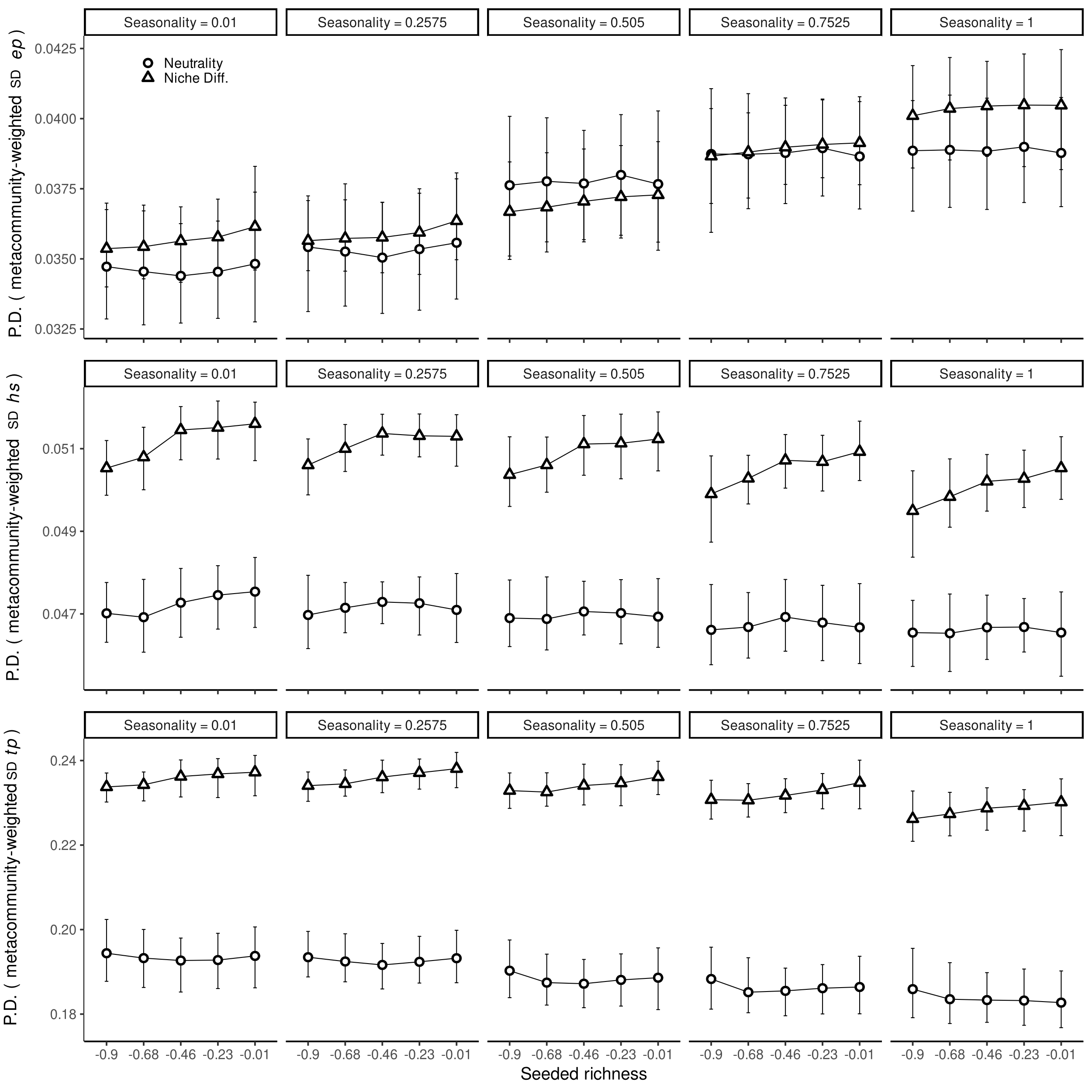

Figure SI. III: Partial dependence (PD) plots showing interactive effects of seasonality (panels), spatial autocorrelation (x-axis), and the niche differentiation assumption (symbols) on the diversity of dispersal strategies in metacommunities (i.e., metacommunity-weighted standard deviation values for ep, hs, ts).

Figure SI.
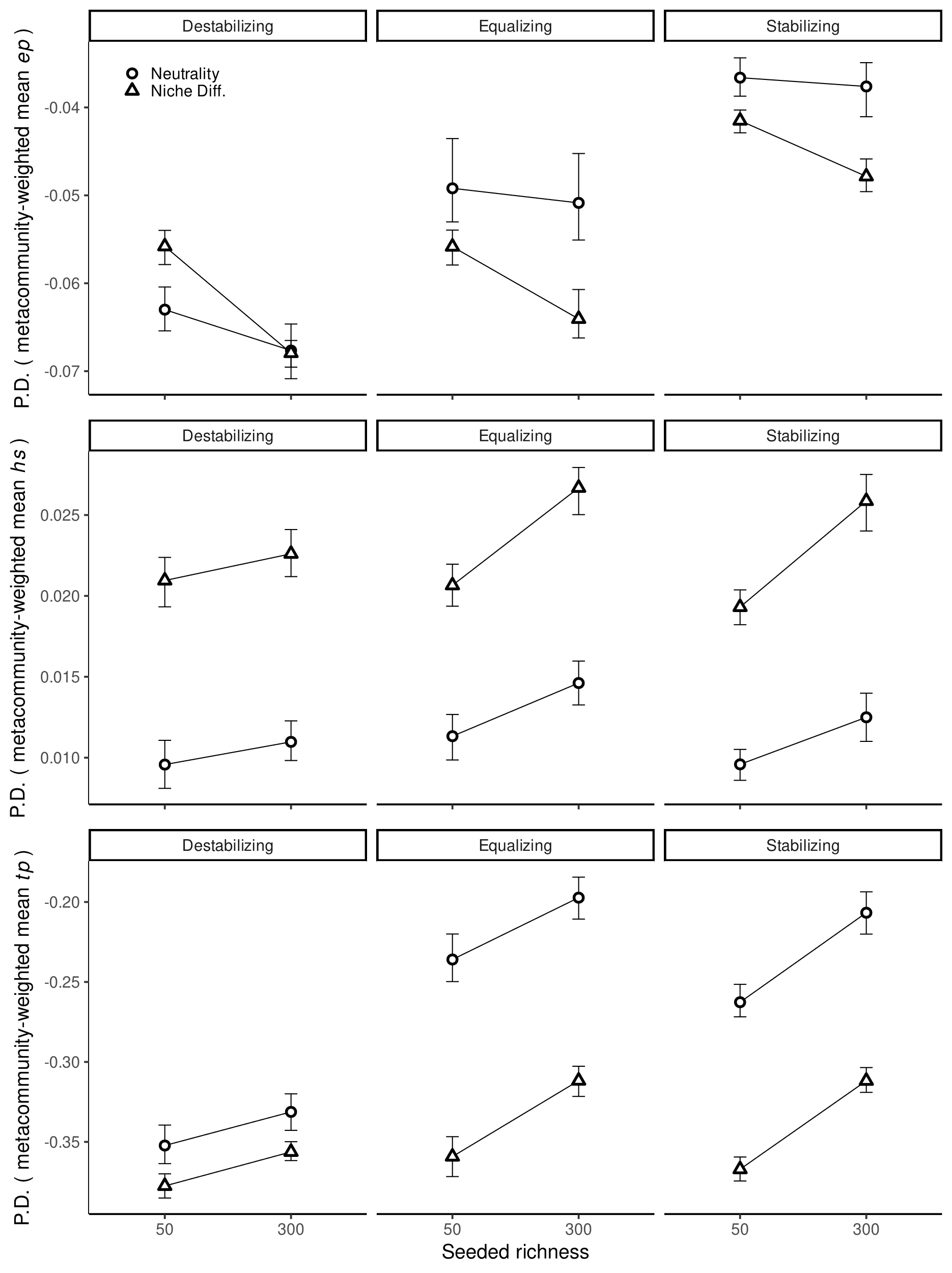
IV: Partial dependence (PD) plots showing interactive effects of competition type (panels), seeded richness of competitors (x-axis), and the niche differentiation assumption (symbols) on the dominant dispersal strategies in metacommunities (i.e., metacommunity-weighted mean values for ep, hs, ts).

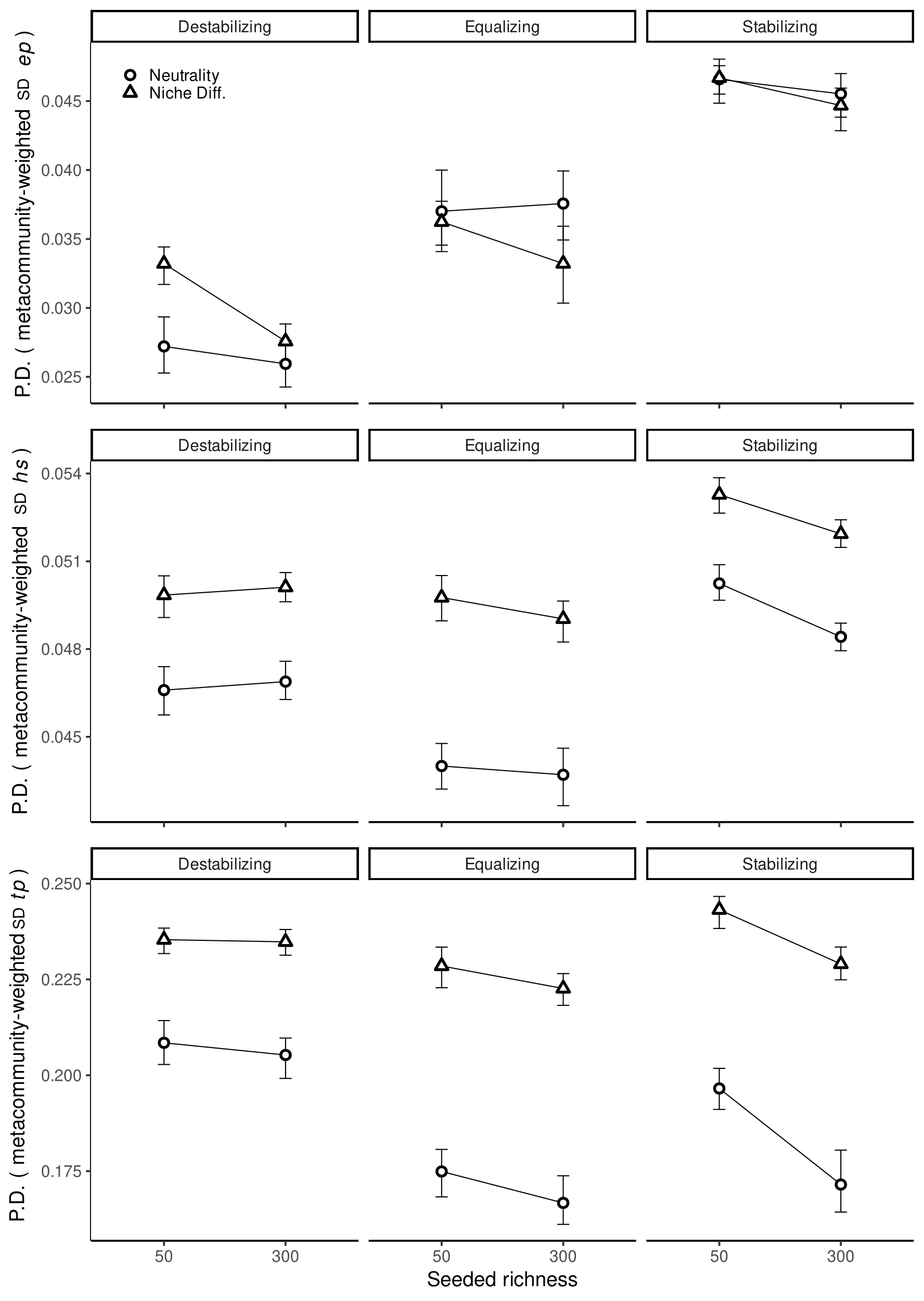

Figure SI.V: Partial dependence (PD) plots showing interactive effects of competition type (panels), seeded richness of competitors (x-axis), and the niche differentiation assumption (symbols) on diversity of dispersal strategies in metacommunities (i.e., metacommunity-weighted standard deviation values for ep, hs, ts).
