## Supplementary material for "Ecological selection of dispersal strategies in metacommunities: impact of landscape features and competitive dynamics": Code and Data to run simulations and generate figures: Analyses and Figures.R

```
library(ggplot2)
library(viridis)
library(WaveletComp)
library(vegan)
library(parallel)
library(reshape2)
library(matrixStats)
library(ape)
library(ggpubr)
library(Boruta)
library(viridis)
library(lattice)
library(gridExtra)
library(randomForest)
library(pdp)
library(randomForestSRC)
library(dplyr)


#Analyses carried out in the manuscript: Ecological selection of dispersal strategies in metacommunities: impact of landscape features and competitive dynamics


#Importing and working the data to generate the figures and run the analyses reported in the main manuscript and SI.
# This dataset presents model summaries across the 20 replicates for simulation scenarios
Data_simulations_final<-read.table("Data_Simulations_Final.txt",sep="\t", dec=".",header = T)


#Subseting the data considering Niche Differentiation
Data_NicheDIFF_Destabilizing_lowdiv<-subset(Data_simulations_final, Niche_diff == "Niche Diff." & N_species==50 & Comp_type=="Destabilizing")
Data_NicheDIFF_Destabilizing_highdiv<-subset(Data_simulations_final, Niche_diff == "Niche Diff." & N_species==300 & Comp_type=="Destabilizing")

Data_NicheDIFF_Equalizing_lowdiv<-subset(Data_simulations_final, Niche_diff == "Niche Diff." & N_species==50 & Comp_type=="Equalizing")
Data_NicheDIFF_Equalizing_highdiv<-subset(Data_simulations_final, Niche_diff == "Niche Diff." & N_species==300 & Comp_type=="Equalizing")

Data_NicheDIFF_Stabilizing_lowdiv<-subset(Data_simulations_final, Niche_diff == "Niche Diff." & N_species==50 & Comp_type=="Stabilizing")
Data_NicheDIFF_Stabilizing_highdiv<-subset(Data_simulations_final, Niche_diff == "Niche Diff." & N_species==300 & Comp_type=="Stabilizing")


Data_simulations_final_Nich_Diff<-subset(Data_simulations_final,Data_simulations_final$Niche_diff=="Niche Diff.")


####Figure 2####

#Heat maps for emigration propensity

Heat_map_Destabilizing_low_div_EP<-dcast(Data_NicheDIFF_Destabilizing_lowdiv[,c("Env_struct","Seasonality","Avg.Weighted_EP")], Env_struct ~Seasonality, value.var="Avg.Weighted_EP")
rownames(Heat_map_Destabilizing_low_div_EP)<-Heat_map_Destabilizing_low_div_EP[,1];Heat_map_Destabilizing_low_div_EP<-Heat_map_Destabilizing_low_div_EP[,-1]
Heat_map_Destabilizing_low_div_EP<-Heat_map_Destabilizing_low_div_EP[c("-0.9","-0.68","-0.46","-0.23","-0.01"),]

Heat_map_equalizing_low_div_EP<-dcast(Data_NicheDIFF_Equalizing_lowdiv[,c("Env_struct","Seasonality","Avg.Weighted_EP")], Env_struct ~Seasonality, value.var="Avg.Weighted_EP")
rownames(Heat_map_equalizing_low_div_EP)<-Heat_map_equalizing_low_div_EP[,1];Heat_map_equalizing_low_div_EP<-Heat_map_equalizing_low_div_EP[,-1]
Heat_map_equalizing_low_div_EP<-Heat_map_equalizing_low_div_EP[c("-0.9","-0.68","-0.46","-0.23","-0.01"),]

Heat_map_stabilizing_low_div_EP<-dcast(Data_NicheDIFF_Stabilizing_lowdiv[,c("Env_struct","Seasonality","Avg.Weighted_EP")], Env_struct ~Seasonality, value.var="Avg.Weighted_EP")
rownames(Heat_map_stabilizing_low_div_EP)<-Heat_map_stabilizing_low_div_EP[,1];Heat_map_stabilizing_low_div_EP<-Heat_map_stabilizing_low_div_EP[,-1]
Heat_map_stabilizing_low_div_EP<-Heat_map_stabilizing_low_div_EP[c("-0.9","-0.68","-0.46","-0.23","-0.01"),]


Heat_map_Destabilizing_high_div_EP<-dcast(Data_NicheDIFF_Destabilizing_highdiv[,c("Env_struct","Seasonality","Avg.Weighted_EP")], Env_struct ~Seasonality, value.var="Avg.Weighted_EP")
rownames(Heat_map_Destabilizing_high_div_EP)<-Heat_map_Destabilizing_high_div_EP[,1];Heat_map_Destabilizing_high_div_EP<-Heat_map_Destabilizing_high_div_EP[,-1]
Heat_map_Destabilizing_high_div_EP<-Heat_map_Destabilizing_high_div_EP[c("-0.9","-0.68","-0.46","-0.23","-0.01"),]


Heat_map_equalizing_high_div_EP<-dcast(Data_NicheDIFF_Equalizing_highdiv[,c("Env_struct","Seasonality","Avg.Weighted_EP")], Env_struct ~Seasonality, value.var="Avg.Weighted_EP")
rownames(Heat_map_equalizing_high_div_EP)<-Heat_map_equalizing_high_div_EP[,1];Heat_map_equalizing_high_div_EP<-Heat_map_equalizing_high_div_EP[,-1]
Heat_map_equalizing_high_div_EP<-Heat_map_equalizing_high_div_EP[c("-0.9","-0.68","-0.46","-0.23","-0.01"),]


Heat_map_stabilizing_high_div_EP<-dcast(Data_NicheDIFF_Stabilizing_highdiv[,c("Env_struct","Seasonality","Avg.Weighted_EP")], Env_struct ~Seasonality, value.var="Avg.Weighted_EP")
rownames(Heat_map_stabilizing_high_div_EP)<-Heat_map_stabilizing_high_div_EP[,1];Heat_map_stabilizing_high_div_EP<-Heat_map_stabilizing_high_div_EP[,-1]
Heat_map_stabilizing_high_div_EP<-Heat_map_stabilizing_high_div_EP[c("-0.9","-0.68","-0.46","-0.23","-0.01"),]

coul=plasma(100)
my.min <- min(unlist(c(Heat_map_Destabilizing_low_div_EP,
                       Heat_map_equalizing_low_div_EP,
                       Heat_map_stabilizing_low_div_EP,
                       Heat_map_Destabilizing_high_div_EP,
                       Heat_map_equalizing_high_div_EP,
                       Heat_map_stabilizing_high_div_EP)))

my.max <- max(unlist(c(Heat_map_Destabilizing_low_div_EP,
                       Heat_map_equalizing_low_div_EP,
                       Heat_map_stabilizing_low_div_EP,
                       Heat_map_Destabilizing_high_div_EP,
                       Heat_map_equalizing_high_div_EP,
                       Heat_map_stabilizing_high_div_EP)))

my.at <- seq(my.min, my.max, length.out=length(coul))
my.ckey <- list(at=my.at, col=coul)

Plot_heat_Dest_low_EP<-levelplot(as.matrix(Heat_map_Destabilizing_low_div_EP),col.regions=coul,at=my.at, colorkey=my.ckey,xlab="", ylab="",scales = list(tck = c(1,0),cex=1.5))
Plot_heat_Equa_low_EP<-levelplot(as.matrix(Heat_map_equalizing_low_div_EP),col.regions=coul,at=my.at, colorkey=my.ckey,xlab="", ylab="",scales = list(tck = c(1,0),cex=1.5))
Plot_heat_Stab_low_EP<-levelplot(as.matrix(Heat_map_stabilizing_low_div_EP),col.regions=coul,at=my.at, colorkey=my.ckey,xlab="", ylab="",scales = list(tck = c(1,0),cex=1.5))

Plot_heat_Dest_high_EP<-levelplot(as.matrix(Heat_map_Destabilizing_high_div_EP),col.regions=coul,at=my.at, colorkey=my.ckey,xlab="", ylab="",scales = list(tck = c(1,0),cex=1.5))
Plot_heat_Equa_high_EP<-levelplot(as.matrix(Heat_map_equalizing_high_div_EP),col.regions=coul,at=my.at, colorkey=my.ckey,xlab="", ylab="",scales = list(tck = c(1,0),cex=1.5))
Plot_heat_Stab_high_EP<-levelplot(as.matrix(Heat_map_stabilizing_high_div_EP),col.regions=coul,at=my.at, colorkey=my.ckey,xlab="", ylab="",scales = list(tck = c(1,0),cex=1.5))

#Heat maps for habitat selection

Heat_map_Destabilizing_low_div_HS<-dcast(Data_NicheDIFF_Destabilizing_lowdiv[,c("Env_struct","Seasonality","Avg.Weighted_HS")], Env_struct ~Seasonality, value.var="Avg.Weighted_HS")
rownames(Heat_map_Destabilizing_low_div_HS)<-Heat_map_Destabilizing_low_div_HS[,1];Heat_map_Destabilizing_low_div_HS<-Heat_map_Destabilizing_low_div_HS[,-1]
Heat_map_Destabilizing_low_div_HS<-Heat_map_Destabilizing_low_div_HS[c("-0.9","-0.68","-0.46","-0.23","-0.01"),]

Heat_map_equalizing_low_div_HS<-dcast(Data_NicheDIFF_Equalizing_lowdiv[,c("Env_struct","Seasonality","Avg.Weighted_HS")], Env_struct ~Seasonality, value.var="Avg.Weighted_HS")
rownames(Heat_map_equalizing_low_div_HS)<-Heat_map_equalizing_low_div_HS[,1];Heat_map_equalizing_low_div_HS<-Heat_map_equalizing_low_div_HS[,-1]
Heat_map_equalizing_low_div_HS<-Heat_map_equalizing_low_div_HS[c("-0.9","-0.68","-0.46","-0.23","-0.01"),]

Heat_map_stabilizing_low_div_HS<-dcast(Data_NicheDIFF_Stabilizing_lowdiv[,c("Env_struct","Seasonality","Avg.Weighted_HS")], Env_struct ~Seasonality, value.var="Avg.Weighted_HS")
rownames(Heat_map_stabilizing_low_div_HS)<-Heat_map_stabilizing_low_div_HS[,1];Heat_map_stabilizing_low_div_HS<-Heat_map_stabilizing_low_div_HS[,-1]
Heat_map_stabilizing_low_div_HS<-Heat_map_stabilizing_low_div_HS[c("-0.9","-0.68","-0.46","-0.23","-0.01"),]


Heat_map_Destabilizing_high_div_HS<-dcast(Data_NicheDIFF_Destabilizing_highdiv[,c("Env_struct","Seasonality","Avg.Weighted_HS")], Env_struct ~Seasonality, value.var="Avg.Weighted_HS")
rownames(Heat_map_Destabilizing_high_div_HS)<-Heat_map_Destabilizing_high_div_HS[,1];Heat_map_Destabilizing_high_div_HS<-Heat_map_Destabilizing_high_div_HS[,-1]
Heat_map_Destabilizing_high_div_HS<-Heat_map_Destabilizing_high_div_HS[c("-0.9","-0.68","-0.46","-0.23","-0.01"),]


Heat_map_equalizing_high_div_HS<-dcast(Data_NicheDIFF_Equalizing_highdiv[,c("Env_struct","Seasonality","Avg.Weighted_HS")], Env_struct ~Seasonality, value.var="Avg.Weighted_HS")
rownames(Heat_map_equalizing_high_div_HS)<-Heat_map_equalizing_high_div_HS[,1];Heat_map_equalizing_high_div_HS<-Heat_map_equalizing_high_div_HS[,-1]
Heat_map_equalizing_high_div_HS<-Heat_map_equalizing_high_div_HS[c("-0.9","-0.68","-0.46","-0.23","-0.01"),]


Heat_map_stabilizing_high_div_HS<-dcast(Data_NicheDIFF_Stabilizing_highdiv[,c("Env_struct","Seasonality","Avg.Weighted_HS")], Env_struct ~Seasonality, value.var="Avg.Weighted_HS")
rownames(Heat_map_stabilizing_high_div_HS)<-Heat_map_stabilizing_high_div_HS[,1];Heat_map_stabilizing_high_div_HS<-Heat_map_stabilizing_high_div_HS[,-1]
Heat_map_stabilizing_high_div_HS<-Heat_map_stabilizing_high_div_HS[c("-0.9","-0.68","-0.46","-0.23","-0.01"),]


coul=mako(100)
my.min <- min(unlist(c(Heat_map_Destabilizing_low_div_HS,
                       Heat_map_equalizing_low_div_HS,
                       Heat_map_stabilizing_low_div_HS,
                       Heat_map_Destabilizing_high_div_HS,
                       Heat_map_equalizing_high_div_HS,
                       Heat_map_stabilizing_high_div_HS)))

my.max <- max(unlist(c(Heat_map_Destabilizing_low_div_HS,
                       Heat_map_equalizing_low_div_HS,
                       Heat_map_stabilizing_low_div_HS,
                       Heat_map_Destabilizing_high_div_HS,
                       Heat_map_equalizing_high_div_HS,
                       Heat_map_stabilizing_high_div_HS)))

my.at <- seq(my.min, my.max, length.out=length(coul))
my.ckey <- list(at=my.at, col=coul)


Plot_heat_Dest_low_HS<-levelplot(as.matrix(Heat_map_Destabilizing_low_div_HS),col.regions=coul,at=my.at, colorkey=my.ckey,xlab="", ylab="",scales = list(tck = c(1,0),cex=1.5))
Plot_heat_Equa_low_HS<-levelplot(as.matrix(Heat_map_equalizing_low_div_HS),col.regions=coul,at=my.at, colorkey=my.ckey,xlab="", ylab="",scales = list(tck = c(1,0),cex=1.5))
Plot_heat_Stab_low_HS<-levelplot(as.matrix(Heat_map_stabilizing_low_div_HS),col.regions=coul,at=my.at, colorkey=my.ckey,xlab="", ylab="",scales = list(tck = c(1,0),cex=1.5))

Plot_heat_Dest_high_HS<-levelplot(as.matrix(Heat_map_Destabilizing_high_div_HS),col.regions=coul,at=my.at, colorkey=my.ckey,xlab="", ylab="",scales = list(tck = c(1,0),cex=1.5))
Plot_heat_Equa_high_HS<-levelplot(as.matrix(Heat_map_equalizing_high_div_HS),col.regions=coul,at=my.at, colorkey=my.ckey,xlab="", ylab="",scales = list(tck = c(1,0),cex=1.5))
Plot_heat_Stab_high_HS<-levelplot(as.matrix(Heat_map_stabilizing_high_div_HS),col.regions=coul,at=my.at, colorkey=my.ckey,xlab="", ylab="",scales = list(tck = c(1,0),cex=1.5))

#Heat maps for traversal capacity

Heat_map_Destabilizing_low_div_TP<-dcast(Data_NicheDIFF_Destabilizing_lowdiv[,c("Env_struct","Seasonality","Avg.Weighted_TP")], Env_struct ~Seasonality, value.var="Avg.Weighted_TP")
rownames(Heat_map_Destabilizing_low_div_TP)<-Heat_map_Destabilizing_low_div_TP[,1];Heat_map_Destabilizing_low_div_TP<-Heat_map_Destabilizing_low_div_TP[,-1]
Heat_map_Destabilizing_low_div_TP<-Heat_map_Destabilizing_low_div_TP[c("-0.9","-0.68","-0.46","-0.23","-0.01"),]

Heat_map_equalizing_low_div_TP<-dcast(Data_NicheDIFF_Equalizing_lowdiv[,c("Env_struct","Seasonality","Avg.Weighted_TP")], Env_struct ~Seasonality, value.var="Avg.Weighted_TP")
rownames(Heat_map_equalizing_low_div_TP)<-Heat_map_equalizing_low_div_TP[,1];Heat_map_equalizing_low_div_TP<-Heat_map_equalizing_low_div_TP[,-1]
Heat_map_equalizing_low_div_TP<-Heat_map_equalizing_low_div_TP[c("-0.9","-0.68","-0.46","-0.23","-0.01"),]

Heat_map_stabilizing_low_div_TP<-dcast(Data_NicheDIFF_Stabilizing_lowdiv[,c("Env_struct","Seasonality","Avg.Weighted_TP")], Env_struct ~Seasonality, value.var="Avg.Weighted_TP")
rownames(Heat_map_stabilizing_low_div_TP)<-Heat_map_stabilizing_low_div_TP[,1];Heat_map_stabilizing_low_div_TP<-Heat_map_stabilizing_low_div_TP[,-1]
Heat_map_stabilizing_low_div_TP<-Heat_map_stabilizing_low_div_TP[c("-0.9","-0.68","-0.46","-0.23","-0.01"),]


Heat_map_Destabilizing_high_div_TP<-dcast(Data_NicheDIFF_Destabilizing_highdiv[,c("Env_struct","Seasonality","Avg.Weighted_TP")], Env_struct ~Seasonality, value.var="Avg.Weighted_TP")
rownames(Heat_map_Destabilizing_high_div_TP)<-Heat_map_Destabilizing_high_div_TP[,1];Heat_map_Destabilizing_high_div_TP<-Heat_map_Destabilizing_high_div_TP[,-1]
Heat_map_Destabilizing_high_div_TP<-Heat_map_Destabilizing_high_div_TP[c("-0.9","-0.68","-0.46","-0.23","-0.01"),]


Heat_map_equalizing_high_div_TP<-dcast(Data_NicheDIFF_Equalizing_highdiv[,c("Env_struct","Seasonality","Avg.Weighted_TP")], Env_struct ~Seasonality, value.var="Avg.Weighted_TP")
rownames(Heat_map_equalizing_high_div_TP)<-Heat_map_equalizing_high_div_TP[,1];Heat_map_equalizing_high_div_TP<-Heat_map_equalizing_high_div_TP[,-1]
Heat_map_equalizing_high_div_TP<-Heat_map_equalizing_high_div_TP[c("-0.9","-0.68","-0.46","-0.23","-0.01"),]


Heat_map_stabilizing_high_div_TP<-dcast(Data_NicheDIFF_Stabilizing_highdiv[,c("Env_struct","Seasonality","Avg.Weighted_TP")], Env_struct ~Seasonality, value.var="Avg.Weighted_TP")
rownames(Heat_map_stabilizing_high_div_TP)<-Heat_map_stabilizing_high_div_TP[,1];Heat_map_stabilizing_high_div_TP<-Heat_map_stabilizing_high_div_TP[,-1]
Heat_map_stabilizing_high_div_TP<-Heat_map_stabilizing_high_div_TP[c("-0.9","-0.68","-0.46","-0.23","-0.01"),]


coul=viridis(100)
my.min <- min(unlist(c(Heat_map_Destabilizing_low_div_TP,
                       Heat_map_equalizing_low_div_TP,
                       Heat_map_stabilizing_low_div_TP,
                       Heat_map_Destabilizing_high_div_TP,
                       Heat_map_equalizing_high_div_TP,
                       Heat_map_stabilizing_high_div_TP)))

my.max <- max(unlist(c(Heat_map_Destabilizing_low_div_TP,
                       Heat_map_equalizing_low_div_TP,
                       Heat_map_stabilizing_low_div_TP,
                       Heat_map_Destabilizing_high_div_TP,
                       Heat_map_equalizing_high_div_TP,
                       Heat_map_stabilizing_high_div_TP)))

my.at <- seq(my.min, my.max, length.out=length(coul))
my.ckey <- list(at=my.at, col=coul)


Plot_heat_Dest_low_TP<-levelplot(as.matrix(Heat_map_Destabilizing_low_div_TP),col.regions=coul,at=my.at, colorkey=my.ckey,xlab="", ylab="",scales = list(tck = c(1,0),cex=1.5))
Plot_heat_Equa_low_TP<-levelplot(as.matrix(Heat_map_equalizing_low_div_TP),col.regions=coul,at=my.at, colorkey=my.ckey,xlab="", ylab="",scales = list(tck = c(1,0),cex=1.5))
Plot_heat_Stab_low_TP<-levelplot(as.matrix(Heat_map_stabilizing_low_div_TP),col.regions=coul,at=my.at, colorkey=my.ckey,xlab="", ylab="",scales = list(tck = c(1,0),cex=1.5))

Plot_heat_Dest_high_TP<-levelplot(as.matrix(Heat_map_Destabilizing_high_div_TP),col.regions=coul,at=my.at, colorkey=my.ckey,xlab="", ylab="",scales = list(tck = c(1,0),cex=1.5))
Plot_heat_Equa_high_TP<-levelplot(as.matrix(Heat_map_equalizing_high_div_TP),col.regions=coul,at=my.at, colorkey=my.ckey,xlab="", ylab="",scales = list(tck = c(1,0),cex=1.5))
Plot_heat_Stab_high_TP<-levelplot(as.matrix(Heat_map_stabilizing_high_div_TP),col.regions=coul,at=my.at, colorkey=my.ckey,xlab="", ylab="",scales = list(tck = c(1,0),cex=1.5))

###### Random forest models#####


#Function to estimate partial dependence and bootstrap CI
partial_plots_rfscr_one_covariate2<-function(formula,Focal_Predictor,covariate,data,repp){
  resample_matrix<-replicate(n = repp,sample(1:nrow(data),replace = T))
  
  if(is.null(covariate)==F){
    
    Matrix_results<-matrix(NA,nrow=length(unique(data[,Focal_Predictor]))*length(unique(data[,covariate])),ncol=repp)
    
    
    
    for(i in 1:ncol(resample_matrix)){
      
      new_data<-data[resample_matrix[,i],]
      Predictors_pairwise<-as.matrix(c(Focal_Predictor,covariate))
      
      model<-randomForest(formula, data = new_data)
      Plot_two_way<-pdp::partial(model,pred.var=Predictors_pairwise[,1],pred.data=new_data)
      Matrix_results[,i]<-Plot_two_way$yhat
      
    }
    
    
    Mean_PD<-rowMeans(Matrix_results)
    
    LIC_PD<-apply(Matrix_results,1,function(xx)quantile(xx,c(0.025)))                            
    
    UIC_PD<-apply(Matrix_results,1,function(xx)quantile(xx,c(0.975)))                            
    
    Data_plot_PD<-data.frame(Plot_two_way[,c(1,2)],Mean_PD,LIC_PD,UIC_PD)
    names(Data_plot_PD)<-c(Focal_Predictor,covariate,"Mean","LIC","UIC")
    
  }else{
    
    
    Matrix_results<-matrix(NA,nrow=length(unique(data[,Focal_Predictor])),ncol=repp)
    
    for(i in 1:ncol(resample_matrix)){
      
      new_data<-data[resample_matrix[,i],]
      Predictors_alone<-as.matrix(c(Focal_Predictor))
      
      model<-randomForest(formula, data = new_data)
      Plot_one_way<-partial(model,pred.var=Predictors_alone[,1],pred.data=new_data)
      Matrix_results[,i]<-Plot_one_way$yhat
    }
    
    
    Mean_PD<-rowMeans(Matrix_results)
    
    LIC_PD<-apply(Matrix_results,1,function(xx)quantile(xx,c(0.025)))                            
    
    UIC_PD<-apply(Matrix_results,1,function(xx)quantile(xx,c(0.975)))                            
    
    Data_plot_PD<-data.frame(Plot_one_way[,1],Mean_PD,LIC_PD,UIC_PD)
    names(Data_plot_PD)<-c(Focal_Predictor,"Mean","LIC","UIC")
    
    
    
  }
  return(Data_plot_PD) 
}


Data_simulations_final$N_species<-factor(Data_simulations_final$N_species,order = TRUE,levels=c("50","300"))

Data_simulations_final$Seasonality<-factor(Data_simulations_final$Seasonality,order = TRUE,levels=unique(Data_simulations_final$Seasonality))
Data_simulations_final$Env_struct<-factor(Data_simulations_final$Env_struct,order = TRUE)
Data_simulations_final$Comp_type<-factor(Data_simulations_final$Comp_type)
Data_simulations_final$Niche_diff<-factor(Data_simulations_final$Niche_diff)

####Metacommunity weighted Mean####


X_train = as.data.frame(Data_simulations_final[,c("N_species","Env_struct","Seasonality","Comp_type","Niche_diff")])
Y_train_TP = Data_simulations_final[,"Avg.Weighted_TP"]
Y_train_HS = Data_simulations_final[,"Avg.Weighted_HS"]
Y_train_EP = Data_simulations_final[,"Avg.Weighted_EP"]


#Feature Selection
boruta_test_TP<- Boruta(Y_train_TP~X_train,data =X_train , doTrace = 2,mcAdj=T,maxRuns=1000)
boruta_test_TP$finalDecision#All confirmed

boruta_test_HS<- Boruta(Y_train_HS~X_train, doTrace = 2,mcAdj=T,maxRuns=1000)
boruta_test_HS$finalDecision# All confirmed

boruta_test_EP<- Boruta(Y_train_EP~X_train, doTrace = 2,mcAdj=T,maxRuns=1000)
boruta_test_EP$finalDecision# Rejected Env.Struct

#Random Forest Models
fit_TP<-rfsrc(Avg.Weighted_TP ~ N_species+Env_struct+Seasonality+Comp_type+Niche_diff, data = Data_simulations_final,importance = "permute")
fit_HS<-rfsrc(Avg.Weighted_HS ~ N_species+Env_struct+Seasonality+Comp_type+Niche_diff, data = Data_simulations_final,importance = "permute")
fit_EP<-rfsrc(Avg.Weighted_EP ~ N_species+Env_struct+Seasonality+Comp_type+Niche_diff, data = Data_simulations_final,importance = "permute")

formula_fit_TP<-formula(Avg.Weighted_TP ~ N_species+Env_struct+Seasonality+Comp_type+Niche_diff)
formula_fit_HS<-formula(Avg.Weighted_HS ~ N_species+Env_struct+Seasonality+Comp_type+Niche_diff)
formula_fit_EP<-formula(Avg.Weighted_EP ~ N_species+Env_struct+Seasonality+Comp_type+Niche_diff)


Test<-randomForest(Avg.Weighted_TP ~ N_species+Env_struct+Seasonality+Comp_type+Niche_diff, data = Data_simulations_final)
#partialPlot(Test,x.var=Env_struct,pred.data=Data_simulations_final)


partial(Test,pred.var=c("Env_struct","N_species","Niche_diff"),pred.data=Data_simulations_final,plot=T)

Plot_two_way<-partial(Test,pred.var=c("Env_struct","Comp_type","Niche_diff"),pred.data=Data_simulations_final)

Predictors_names<-c("N_species","Env_struct","Seasonality","Comp_type")


options(rf.cores = 1)

start_time_full<-Sys.time() 

#Bootstraping
TPPDs<-lapply(1:length(Predictors_names),function(ppp)
  partial_plots_rfscr_one_covariate2(formula=formula_fit_TP,Focal_Predictor=Predictors_names[ppp],covariate ="Niche_diff",data=Data_simulations_final,repp=300))


HSPDs<-lapply(1:length(Predictors_names),function(ppp)
  partial_plots_rfscr_one_covariate2(formula=formula_fit_HS,Focal_Predictor=Predictors_names[ppp],covariate ="Niche_diff",data=Data_simulations_final,repp=300))

EPPDs<-lapply(1:length(Predictors_names),function(ppp)
  partial_plots_rfscr_one_covariate2(formula=formula_fit_EP,Focal_Predictor=Predictors_names[ppp],covariate ="Niche_diff",data=Data_simulations_final,repp=300))

end_time_full<-Sys.time()

end_time_full-start_time_full


Min_Max_HS<-c(min(sapply(HSPDs,function(x)min(x$LIC))),
              max(sapply(HSPDs,function(x)max(x$UIC))))

Min_Max_EP<-c(min(sapply(EPPDs,function(x)min(x$LIC))),
              max(sapply(EPPDs,function(x)max(x$UIC))))

Min_Max_TP<-c(min(sapply(TPPDs,function(x)min(x$LIC))),
               max(sapply(TPPDs,function(x)max(x$UIC))))


#Spatial struct plot##

HS_Spatial_strucutre<-ggplot(HSPDs[[2]],aes(x=Env_struct,y=Mean,group=Niche_diff))+
  ylim(Min_Max_HS)+
  geom_errorbar(aes(ymin=LIC,ymax=UIC),width=.1)+
  geom_line(data = HSPDs[[2]],
            mapping = aes(x = Env_struct, y = Mean, group=Niche_diff))+
  ylab(expression('P.D. ( metacommunity-weighted mean'~italic(hs)~")"))+
  xlab("Spatial autocorrelation")+
  geom_point(aes(shape=Niche_diff),size=8,alpha=1,fill="white",stroke=2)+
  scale_shape_manual(values=c(21, 24))+  theme_classic(base_size = 30)+
  theme(legend.position = "none")

TP_Spatial_strucutre<-ggplot(TPPDs[[2]],aes(x=Env_struct,y=Mean,group=Niche_diff))+
  ylim(Min_Max_TP)+
  geom_errorbar(aes(ymin=LIC,ymax=UIC),width=.1)+
  geom_line(data = TPPDs[[2]],
            mapping = aes(x = Env_struct, y = Mean, group=Niche_diff))+
  ylab(expression('P.D. ( metacommunity-weighted mean'~italic(tp)~")"))+
  xlab("Spatial autocorrelation")+
  geom_point(aes(shape=Niche_diff),size=8,alpha=1,fill="white",stroke=2)+
  scale_shape_manual(values=c(21, 24))+  theme_classic(base_size = 30)+
  theme(legend.position = "none")


EP_Spatial_strucutre<-ggplot(EPPDs[[2]],aes(x=Env_struct,y=Mean,group=Niche_diff))+
  ylim(Min_Max_EP)+
  geom_errorbar(aes(ymin=LIC,ymax=UIC),width=.1)+
  geom_line(data = EPPDs[[2]],
            mapping = aes(x = Env_struct, y = Mean, group=Niche_diff))+
  ylab(expression('P.D. ( metacommunity-weighted mean'~italic(ep)~")"))+
  xlab("Spatial autocorrelation")+
  geom_point(aes(shape=Niche_diff),size=8,alpha=1,fill="white",stroke=2)+
  scale_shape_manual(values=c(21, 24))+  theme_classic(base_size = 30)+
  theme(legend.position = "none")

#Seasonality plots plot##

HS_Seasonality<-ggplot(HSPDs[[3]],aes(x=Seasonality,y=Mean,group=Niche_diff))+
  ylim(Min_Max_HS)+
  geom_errorbar(aes(ymin=LIC,ymax=UIC),width=.1)+
  geom_line(data = HSPDs[[3]],
            mapping = aes(x = Seasonality, y = Mean, group=Niche_diff))+
  ylab(expression('P.D. ( metacommunity-weighted mean'~italic(hs)~")"))+
  xlab("Seasonality")+
  geom_point(aes(shape=Niche_diff),size=8,alpha=1,fill="white",stroke=2)+
  scale_shape_manual(values=c(21, 24))+  theme_classic(base_size = 30)+
  theme(legend.position = "none")

TP_Seasonality<-ggplot(TPPDs[[3]],aes(x=Seasonality,y=Mean,group=Niche_diff))+
  ylim(Min_Max_TP)+
  geom_errorbar(aes(ymin=LIC,ymax=UIC),width=.1)+
  geom_line(data = TPPDs[[3]],
            mapping = aes(x = Seasonality, y = Mean, group=Niche_diff))+
  ylab(expression('P.D. ( metacommunity-weighted mean'~italic(tp)~")"))+
  xlab("Seasonality")+
  geom_point(aes(shape=Niche_diff),size=8,alpha=1,fill="white",stroke=2)+
  scale_shape_manual(values=c(21, 24))+  theme_classic(base_size = 30)+
  theme(legend.position = "none")


EP_Seasonality<-ggplot(EPPDs[[3]],aes(x=Seasonality,y=Mean,group=Niche_diff))+
  ylim(Min_Max_EP)+
  geom_errorbar(aes(ymin=LIC,ymax=UIC),width=.1)+
  geom_line(data = EPPDs[[3]],
            mapping = aes(x = Seasonality, y = Mean, group=Niche_diff))+
  ylab(expression('P.D. ( metacommunity-weighted mean'~italic(ep)~")"))+
  xlab("Seasonality")+
  geom_point(aes(shape=Niche_diff),size=8,alpha=1,fill="white",stroke=2)+
  scale_shape_manual(values=c(21, 24))+  
  theme_classic(base_size = 30)+
  theme(legend.position = c(0.3, 0.8),
        legend.title=element_blank())

## Richness##
N_Species_HS<-ggplot(HSPDs[[1]],aes(x=N_species,y=Mean,group=Niche_diff))+
  ylim(Min_Max_HS)+
  geom_errorbar(aes(ymin=LIC,ymax=UIC),width=.1)+
  geom_line(data = HSPDs[[1]],
            mapping = aes(x = N_species, y = Mean, group=Niche_diff))+
  ylab(expression('P.D. ( metacommunity-weighted mean'~italic(hs)~")"))+
  xlab("Seeded richness")+
  geom_point(aes(shape=Niche_diff),size=8,alpha=1,fill="white",stroke=2)+
  scale_shape_manual(values=c(21, 24))+  theme_classic(base_size = 30)+
  theme(legend.position = "none")

N_Species_TP<-ggplot(TPPDs[[1]],aes(x=N_species,y=Mean,group=Niche_diff))+
  ylim(Min_Max_TP)+
  geom_errorbar(aes(ymin=LIC,ymax=UIC),width=.1)+
  geom_line(data = TPPDs[[1]],
            mapping = aes(x = N_species, y = Mean, group=Niche_diff))+
  ylab(expression('P.D. ( metacommunity-weighted mean'~italic(tp)~")"))+
  xlab("Seeded richness")+
  geom_point(aes(shape=Niche_diff),size=8,alpha=1,fill="white",stroke=2)+
  scale_shape_manual(values=c(21, 24))+  theme_classic(base_size = 30)+
  theme(legend.position = "none")


N_Species_EP<-ggplot(EPPDs[[1]],aes(x=N_species,y=Mean,group=Niche_diff))+
  ylim(Min_Max_EP)+
  geom_errorbar(aes(ymin=LIC,ymax=UIC),width=.1)+
  geom_line(data = EPPDs[[1]],
            mapping = aes(x = N_species, y = Mean, group=Niche_diff))+
  ylab(expression('P.D. ( metacommunity-weighted mean'~italic(ep)~")"))+
  xlab("Seeded richness")+
  geom_point(aes(shape=Niche_diff),size=8,alpha=1,fill="white",stroke=2)+
  scale_shape_manual(values=c(21, 24))+
  theme_classic(base_size = 20)+
  theme(legend.position = "none")

#Competition type##

Comp_type_HS<-ggplot(HSPDs[[4]],aes(x=Comp_type,y=Mean,group=Niche_diff))+
  ylim(Min_Max_HS)+
  geom_errorbar(aes(ymin=LIC,ymax=UIC),width=.1)+
  ylab(expression('P.D. ( metacommunity-weighted mean'~italic(hs)~")"))+
  xlab("Competition Type")+
  geom_point(aes(shape=Niche_diff),size=8,alpha=1,fill="white",stroke=2)+
  scale_shape_manual(values=c(21, 24))+  theme_classic(base_size = 30)+
  theme(legend.position = "none")

Comp_type_TP<-ggplot(TPPDs[[4]],aes(x=Comp_type,y=Mean,group=Niche_diff))+
  ylim(Min_Max_TP)+
  geom_errorbar(aes(ymin=LIC,ymax=UIC),width=.1)+
  ylab(expression('P.D. ( metacommunity-weighted mean'~italic(tp)~")"))+
  xlab("Competition Type")+
  geom_point(aes(shape=Niche_diff),size=8,alpha=1,fill="white",stroke=2)+
  scale_shape_manual(values=c(21, 24))+  theme_classic(base_size = 30)+
  theme(legend.position = "none")


Comp_type_EP<-ggplot(EPPDs[[4]],aes(x=Comp_type,y=Mean,group=Niche_diff))+
  ylim(Min_Max_EP)+
  geom_errorbar(aes(ymin=LIC,ymax=UIC),width=.1)+
  ylab(expression('P.D. ( metacommunity-weighted mean'~italic(ep)~")"))+
  xlab("Competition Type")+
  geom_point(aes(shape=Niche_diff),size=8,alpha=1,fill="white",stroke=2)+
  scale_shape_manual(values=c(21, 24))+  theme_classic(base_size = 30)+
  theme(legend.position = "none")

#Figure 3
egg::ggarrange(EP_Seasonality+rremove("x.title")+rremove("x.text"),EP_Spatial_strucutre+rremove("xylab")+rremove("xy.text"),N_Species_EP+rremove("xylab")+rremove("xy.text"),Comp_type_EP+rremove("xylab")+rremove("xy.text"),
               TP_Seasonality+rremove("x.title")+rremove("x.text"),TP_Spatial_strucutre+rremove("xylab")+rremove("xy.text"),N_Species_TP+rremove("xylab")+rremove("xy.text"),Comp_type_TP+rremove("xylab")+rremove("xy.text"),
               TP_Seasonality,TP_Spatial_strucutre+rremove("ylab")+rremove("y.text"),N_Species_TP+rremove("ylab")+rremove("y.text"),Comp_type_TP+rremove("ylab")+rremove("y.text"),
               ncol=4,nrow = 3)

#####Metacommunity Weighted SD#####

X_train = as.data.frame(Data_simulations_final[,c("N_species","Env_struct","Seasonality","Comp_type","Niche_diff")])
Y_train_TP_SD = Data_simulations_final[,"SD_TP"]
Y_train_HS_SD = Data_simulations_final[,"SD_HS"]
Y_train_EP_SD = Data_simulations_final[,"SD_EP"]


boruta_test_TP_SD<- Boruta(Y_train_TP_SD~X_train, doTrace = 2,mcAdj=T,maxRuns=1000)
boruta_test_TP_SD$finalDecision#Env Struc rejected, Seasonality Tentative

boruta_test_HS_SD<- Boruta(Y_train_HS_SD~X_train, doTrace = 2,mcAdj=T,maxRuns=1000)
boruta_test_HS_SD$finalDecision# Rejected Seasonality and Env Struc

boruta_test_EP_SD<- Boruta(Y_train_EP_SD~X_train, doTrace = 2,mcAdj=T,maxRuns=1000)
boruta_test_EP_SD$finalDecision# Rejected Env.Struct


fit_TP_SD<-rfsrc(SD_TP ~ N_species+Env_struct+Seasonality+Comp_type+Niche_diff, data = Data_simulations_final,importance = "permute")
fit_HS_SD<-rfsrc(SD_HS ~ N_species+Env_struct+Seasonality+Comp_type+Niche_diff, data = Data_simulations_final,importance = "permute")
fit_EP_SD<-rfsrc(SD_EP ~ N_species+Env_struct+Seasonality+Comp_type+Niche_diff, data = Data_simulations_final,importance = "permute")

formula_fit_TP_SD<-formula(SD_TP ~ N_species+Env_struct+Seasonality+Comp_type+Niche_diff)
formula_fit_HS_SD<-formula(SD_HS ~ N_species+Env_struct+Seasonality+Comp_type+Niche_diff)
formula_fit_EP_SD<-formula(SD_EP ~ N_species+Env_struct+Seasonality+Comp_type+Niche_diff)
#Bootstraping

TPPDs_SD<-lapply(1:length(Predictors_names),function(ppp)
  partial_plots_rfscr_one_covariate2(formula=formula_fit_TP_SD,Focal_Predictor=Predictors_names[ppp],covariate ="Niche_diff",data=Data_simulations_final,repp=300))


HSPDs_SD<-lapply(1:length(Predictors_names),function(ppp)
  partial_plots_rfscr_one_covariate2(formula=formula_fit_HS_SD,Focal_Predictor=Predictors_names[ppp],covariate ="Niche_diff",data=Data_simulations_final,repp=300))

EPPDs_SD<-lapply(1:length(Predictors_names),function(ppp)
  partial_plots_rfscr_one_covariate2(formula=formula_fit_EP_SD,Focal_Predictor=Predictors_names[ppp],covariate ="Niche_diff",data=Data_simulations_final,repp=300))


Min_Max_HS<-c(min(sapply(HSPDs_SD,function(x)min(x$LIC))),
              max(sapply(HSPDs_SD,function(x)max(x$UIC))))

Min_Max_EP<-c(min(sapply(EPPDs_SD,function(x)min(x$LIC))),
              max(sapply(EPPDs_SD,function(x)max(x$UIC))))

Min_Max_TP<-c(min(sapply(TPPDs_SD,function(x)min(x$LIC))),
               max(sapply(TPPDs_SD,function(x)max(x$UIC))))


#Spatial struct plot##

HS_Spatial_strucutre_SD<-ggplot(HSPDs_SD[[2]],aes(x=Env_struct,y=Mean,group=Niche_diff))+
  ylim(Min_Max_HS)+
  geom_errorbar(aes(ymin=LIC,ymax=UIC),width=.1)+
  geom_line(data = HSPDs_SD[[2]],
            mapping = aes(x = Env_struct, y = Mean, group=Niche_diff))+
  ylab(expression('P.D. ( metacommunity-weighted SD'~italic(hs)~")"))+
  xlab("Spatial autocorrelation")+
  geom_point(aes(shape=Niche_diff),size=8,alpha=1,fill="white",stroke=2)+
  scale_shape_manual(values=c(21, 24))+  theme_classic(base_size = 30)+
  theme(legend.position = "none")

TP_Spatial_strucutre_SD<-ggplot(TPPDs_SD[[2]],aes(x=Env_struct,y=Mean,group=Niche_diff))+
  ylim(Min_Max_TP)+
  geom_errorbar(aes(ymin=LIC,ymax=UIC),width=.1)+
  geom_line(data = TPPDs_SD[[2]],
            mapping = aes(x = Env_struct, y = Mean, group=Niche_diff))+
  ylab(expression('P.D. ( metacommunity-weighted SD'~italic(tp)~")"))+
  xlab("Spatial autocorrelation")+
  geom_point(aes(shape=Niche_diff),size=8,alpha=1,fill="white",stroke=2)+
  scale_shape_manual(values=c(21, 24))+  theme_classic(base_size = 30)+
  theme(legend.position = "none")


EP_Spatial_strucutre_SD<-ggplot(EPPDs_SD[[2]],aes(x=Env_struct,y=Mean,group=Niche_diff))+
  ylim(Min_Max_EP)+
  geom_errorbar(aes(ymin=LIC,ymax=UIC),width=.1)+
  geom_line(data = EPPDs_SD[[2]],
            mapping = aes(x = Env_struct, y = Mean, group=Niche_diff))+
  ylab(expression('P.D. ( metacommunity-weighted SD'~italic(ep)~")"))+
  xlab("Spatial autocorrelation")+
  geom_point(aes(shape=Niche_diff),size=8,alpha=1,fill="white",stroke=2)+
  scale_shape_manual(values=c(21, 24))+  theme_classic(base_size = 30)+
  theme(legend.position = "none")

#Seasonality plots plot##

HS_Seasonality_SD<-ggplot(HSPDs_SD[[3]],aes(x=Seasonality,y=Mean,group=Niche_diff))+
  ylim(Min_Max_HS)+
  geom_errorbar(aes(ymin=LIC,ymax=UIC),width=.1)+
  geom_line(data = HSPDs_SD[[3]],
            mapping = aes(x = Seasonality, y = Mean, group=Niche_diff))+
  ylab(expression('P.D. ( metacommunity-weighted SD'~italic(hs)~")"))+
  xlab("Seasonality")+
  geom_point(aes(shape=Niche_diff),size=8,alpha=1,fill="white",stroke=2)+
  scale_shape_manual(values=c(21, 24))+  theme_classic(base_size = 30)+
  theme(legend.position = "none")

TP_Seasonality_SD<-ggplot(TPPDs_SD[[3]],aes(x=Seasonality,y=Mean,group=Niche_diff))+
  ylim(Min_Max_TP)+
  geom_errorbar(aes(ymin=LIC,ymax=UIC),width=.1)+
  geom_line(data = TPPDs_SD[[3]],
            mapping = aes(x = Seasonality, y = Mean, group=Niche_diff))+
  ylab(expression('P.D. ( metacommunity-weighted SD'~italic(tp)~")"))+
  xlab("Seasonality")+
  geom_point(aes(shape=Niche_diff),size=8,alpha=1,fill="white",stroke=2)+
  scale_shape_manual(values=c(21, 24))+  theme_classic(base_size = 30)+
  theme(legend.position = "none")


EP_Seasonality_SD<-ggplot(EPPDs_SD[[3]],aes(x=Seasonality,y=Mean,group=Niche_diff))+
  ylim(Min_Max_EP)+
  geom_errorbar(aes(ymin=LIC,ymax=UIC),width=.1)+
  geom_line(data = EPPDs_SD[[3]],
            mapping = aes(x = Seasonality, y = Mean, group=Niche_diff))+
  ylab(expression('P.D. ( metacommunity-weighted SD'~italic(ep)~")"))+
  xlab("Seasonality")+
  geom_point(aes(shape=Niche_diff),size=8,alpha=1,fill="white",stroke=2)+
  scale_shape_manual(values=c(21, 24))+  
  theme_classic(base_size = 30)+
  theme(legend.position = c(0.3, 0.8),
        legend.title=element_blank())

## Richness##
N_Species_HS_SD<-ggplot(HSPDs_SD[[1]],aes(x=N_species,y=Mean,group=Niche_diff))+
  ylim(Min_Max_HS)+
  geom_errorbar(aes(ymin=LIC,ymax=UIC),width=.1)+
  geom_line(data = HSPDs_SD[[1]],
            mapping = aes(x = N_species, y = Mean, group=Niche_diff))+
  ylab(expression('P.D. ( metacommunity-weighted SD'~italic(hs)~")"))+
  xlab("Seeded richness")+
  geom_point(aes(shape=Niche_diff),size=8,alpha=1,fill="white",stroke=2)+
  scale_shape_manual(values=c(21, 24))+  theme_classic(base_size = 30)+
  theme(legend.position = "none")

N_Species_TP_SD<-ggplot(TPPDs_SD[[1]],aes(x=N_species,y=Mean,group=Niche_diff))+
  ylim(Min_Max_TP)+
  geom_errorbar(aes(ymin=LIC,ymax=UIC),width=.1)+
  geom_line(data = TPPDs_SD[[1]],
            mapping = aes(x = N_species, y = Mean, group=Niche_diff))+
  ylab(expression('P.D. ( metacommunity-weighted SD'~italic(tp)~")"))+
  xlab("Seeded richness")+
  geom_point(aes(shape=Niche_diff),size=8,alpha=1,fill="white",stroke=2)+
  scale_shape_manual(values=c(21, 24))+  theme_classic(base_size = 30)+
  theme(legend.position = "none")


N_Species_EP_SD<-ggplot(EPPDs_SD[[1]],aes(x=N_species,y=Mean,group=Niche_diff))+
  ylim(Min_Max_EP)+
  geom_errorbar(aes(ymin=LIC,ymax=UIC),width=.1)+
  geom_line(data = EPPDs_SD[[1]],
            mapping = aes(x = N_species, y = Mean, group=Niche_diff))+
  ylab(expression('P.D. ( metacommunity-weighted SD'~italic(ep)~")"))+
  xlab("Seeded richness")+
  geom_point(aes(shape=Niche_diff),size=8,alpha=1,fill="white",stroke=2)+
  scale_shape_manual(values=c(21, 24))+
  theme_classic(base_size = 20)+
  theme(legend.position = "none")

#Competition type##

Comp_type_HS_SD<-ggplot(HSPDs_SD[[4]],aes(x=Comp_type,y=Mean,group=Niche_diff))+
  ylim(Min_Max_HS)+
  geom_errorbar(aes(ymin=LIC,ymax=UIC),width=.1)+
  ylab(expression('P.D. ( metacommunity-weighted SD'~italic(hs)~")"))+
  xlab("Competition Type")+
  geom_point(aes(shape=Niche_diff),size=8,alpha=1,fill="white",stroke=2)+
  scale_shape_manual(values=c(21, 24))+  theme_classic(base_size = 30)+
  theme(legend.position = "none")

Comp_type_TP_SD<-ggplot(TPPDs_SD[[4]],aes(x=Comp_type,y=Mean,group=Niche_diff))+
  ylim(Min_Max_TP)+
  geom_errorbar(aes(ymin=LIC,ymax=UIC),width=.1)+
  ylab(expression('P.D. ( metacommunity-weighted SD'~italic(tp)~")"))+
  xlab("Competition Type")+
  geom_point(aes(shape=Niche_diff),size=8,alpha=1,fill="white",stroke=2)+
  scale_shape_manual(values=c(21, 24))+  theme_classic(base_size = 30)+
  theme(legend.position = "none")


Comp_type_EP_SD<-ggplot(EPPDs_SD[[4]],aes(x=Comp_type,y=Mean,group=Niche_diff))+
  ylim(Min_Max_EP)+
  geom_errorbar(aes(ymin=LIC,ymax=UIC),width=.1)+
  ylab(expression('P.D. ( metacommunity-weighted SD'~italic(ep)~")"))+
  xlab("Competition Type")+
  geom_point(aes(shape=Niche_diff),size=8,alpha=1,fill="white",stroke=2)+
  scale_shape_manual(values=c(21, 24))+  theme_classic(base_size = 30)+
  theme(legend.position = "none")

#Figure SI.I
egg::ggarrange(EP_Seasonality_SD+rremove("x.title")+rremove("x.text"),EP_Spatial_strucutre_SD+rremove("xylab")+rremove("xy.text"),N_Species_EP_SD+rremove("xylab")+rremove("xy.text"),Comp_type_EP_SD+rremove("xylab")+rremove("xy.text"),
               HS_Seasonality_SD+rremove("x.title")+rremove("x.text"),HS_Spatial_strucutre_SD+rremove("xylab")+rremove("xy.text"),N_Species_HS_SD+rremove("xylab")+rremove("xy.text"),Comp_type_HS_SD+rremove("xylab")+rremove("xy.text"),
               TP_Seasonality_SD,TP_Spatial_strucutre_SD+rremove("ylab")+rremove("y.text"),N_Species_TP_SD+rremove("ylab")+rremove("y.text"),Comp_type_TP_SD+rremove("ylab")+rremove("y.text"),
               ncol=4,nrow = 3)

##### Supp Information figures (3-way interactions)####

partial_plots_rfscr_one_covariate3<-function(formula,Focal_Predictor,covariate,covariate2,data,repp){
  resample_matrix<-replicate(n = repp,sample(1:nrow(data),replace = T))
  
  if(is.null(covariate)==F){
    
    Matrix_results<-matrix(NA,nrow=length(unique(data[,Focal_Predictor]))*length(unique(data[,covariate]))*length(unique(data[,covariate2])),ncol=repp)
    
    
    
    for(i in 1:ncol(resample_matrix)){
      
      new_data<-data[resample_matrix[,i],]
      Predictors_pairwise<-as.matrix(c(Focal_Predictor,covariate,covariate2))
      
      model<-randomForest(formula, data = new_data)
      Plot_two_way<-pdp::partial(model,pred.var=Predictors_pairwise[,1],pred.data=new_data)
      Matrix_results[,i]<-Plot_two_way$yhat
      
    }
    
    
    Mean_PD<-rowMeans(Matrix_results)
    
    LIC_PD<-apply(Matrix_results,1,function(xx)quantile(xx,c(0.025)))                            
    
    UIC_PD<-apply(Matrix_results,1,function(xx)quantile(xx,c(0.975)))                            
    
    Data_plot_PD<-data.frame(Plot_two_way[,c(1:3)],Mean_PD,LIC_PD,UIC_PD)
    names(Data_plot_PD)<-c(Focal_Predictor,covariate,covariate2,"Mean","LIC","UIC")
    
  }else{
    
    
    Matrix_results<-matrix(NA,nrow=length(unique(data[,Focal_Predictor])),ncol=repp)
    
    for(i in 1:ncol(resample_matrix)){
      
      new_data<-data[resample_matrix[,i],]
      Predictors_alone<-as.matrix(c(Focal_Predictor))
      
      model<-randomForest(formula, data = new_data)
      Plot_one_way<-pdp::partial(model,pred.var=Predictors_alone[,1],pred.data=new_data)
      Matrix_results[,i]<-Plot_one_way$yhat
    }
    
    
    Mean_PD<-rowMeans(Matrix_results)
    
    LIC_PD<-apply(Matrix_results,1,function(xx)quantile(xx,c(0.025)))                            
    
    UIC_PD<-apply(Matrix_results,1,function(xx)quantile(xx,c(0.975)))                            
    
    Data_plot_PD<-data.frame(Plot_one_way[,1],Mean_PD,LIC_PD,UIC_PD)
    names(Data_plot_PD)<-c(Focal_Predictor,"Mean","LIC","UIC")
    
    
    
  }
  return(Data_plot_PD) 
}


#Seasonality * Spatial Struct. * Niche Diff


EP_Seasonality_Spatial_struct<-partial_plots_rfscr_one_covariate3(formula=formula_fit_EP,Focal_Predictor="Env_struct",covariate="Seasonality",covariate2 ="Niche_diff",data=Data_simulations_final,repp=100)
HS_Seasonality_Spatial_struct<-partial_plots_rfscr_one_covariate3(formula=formula_fit_HS,Focal_Predictor="Env_struct",covariate="Seasonality",covariate2 ="Niche_diff",data=Data_simulations_final,repp=100)
TP_Seasonality_Spatial_struct<-partial_plots_rfscr_one_covariate3(formula=formula_fit_TP,Focal_Predictor="Env_struct",covariate="Seasonality",covariate2 ="Niche_diff",data=Data_simulations_final,repp=100)

EP_Seasonality_Spatial_struct_SD<-partial_plots_rfscr_one_covariate3(formula=formula_fit_EP_SD,Focal_Predictor="Env_struct",covariate="Seasonality",covariate2 ="Niche_diff",data=Data_simulations_final,repp=100)
HS_Seasonality_Spatial_struct_SD<-partial_plots_rfscr_one_covariate3(formula=formula_fit_HS_SD,Focal_Predictor="Env_struct",covariate="Seasonality",covariate2 ="Niche_diff",data=Data_simulations_final,repp=100)
TP_Seasonality_Spatial_struct_SD<-partial_plots_rfscr_one_covariate3(formula=formula_fit_TP_SD,Focal_Predictor="Env_struct",covariate="Seasonality",covariate2 ="Niche_diff",data=Data_simulations_final,repp=100)


EP_Seasonality_Spatial_struct$Seasonality2<-as.factor(dplyr::recode(EP_Seasonality_Spatial_struct$Seasonality, 
                                                                    "0.01" ="Seasonality = 0.01", 
                                                                    "0.2575" = "Seasonality = 0.2575",
                                                                    "0.505"= "Seasonality = 0.505",
                                                                    "0.7525"="Seasonality = 0.7525",
                                                                    "1"="Seasonality = 1"))


TP_Seasonality_Spatial_struct$Seasonality2<-as.factor(dplyr::recode(TP_Seasonality_Spatial_struct$Seasonality, 
                                                                    "0.01" ="Seasonality = 0.01", 
                                                                    "0.2575" = "Seasonality = 0.2575",
                                                                    "0.505"= "Seasonality = 0.505",
                                                                    "0.7525"="Seasonality = 0.7525",
                                                                    "1"="Seasonality = 1"))


HS_Seasonality_Spatial_struct$Seasonality2<-as.factor(dplyr::recode(HS_Seasonality_Spatial_struct$Seasonality, 
                                                                     "0.01" ="Seasonality = 0.01", 
                                                                     "0.2575" = "Seasonality = 0.2575",
                                                                     "0.505"= "Seasonality = 0.505",
                                                                     "0.7525"="Seasonality = 0.7525",
                                                                     "1"="Seasonality = 1"))


EP_Seasonality_Spatial_struct_plot<-ggplot(EP_Seasonality_Spatial_struct,aes(x=Env_struct,y=Mean,group=Niche_diff))+
  #ylim(Min_Max_EP)+
  geom_errorbar(aes(ymin=LIC,ymax=UIC),width=.1)+
  geom_line(data = EP_Seasonality_Spatial_struct,
            mapping = aes(x = Env_struct, y = Mean, group=Niche_diff))+
  ylab(expression('P.D. ( metacommunity-weighted mean'~italic(ep)~")"))+
  xlab("Spatial autocorrelation")+
  geom_point(aes(shape=Niche_diff),size=3,alpha=1,fill="white",stroke=2)+
  facet_wrap(~Seasonality2,ncol = 5)+
  scale_shape_manual(values=c(21, 24))+
  theme_classic(base_size = 20)+
  theme(legend.position = c(0.1, 0.9),
        legend.title=element_blank())


HS_Seasonality_Spatial_struct_plot<-ggplot(HS_Seasonality_Spatial_struct,aes(x=Env_struct,y=Mean,group=Niche_diff))+
  #ylim(Min_Max_EP)+
  geom_errorbar(aes(ymin=LIC,ymax=UIC),width=.1)+
  geom_line(data = HS_Seasonality_Spatial_struct,
            mapping = aes(x = Env_struct, y = Mean, group=Niche_diff))+
  ylab(expression('P.D. ( metacommunity-weighted mean'~italic(hs)~")"))+
  xlab("Spatial autocorrelation")+
  geom_point(aes(shape=Niche_diff),size=3,alpha=1,fill="white",stroke=2)+
  facet_wrap(~Seasonality2,ncol = 5)+
  scale_shape_manual(values=c(21, 24))+
  theme_classic(base_size = 20)+
  theme(legend.position = "none")


TP_Seasonality_Spatial_struct_plot<-ggplot(TP_Seasonality_Spatial_struct,aes(x=Env_struct,y=Mean,group=Niche_diff))+
  #ylim(Min_Max_EP)+
  geom_errorbar(aes(ymin=LIC,ymax=UIC),width=.1)+
  geom_line(data = TP_Seasonality_Spatial_struct,
            mapping = aes(x = Env_struct, y = Mean, group=Niche_diff))+
  ylab(expression('P.D. ( metacommunity-weighted mean'~italic(tp)~")"))+
  xlab("Spatial autocorrelation")+
  geom_point(aes(shape=Niche_diff),size=3,alpha=1,fill="white",stroke=2)+
  facet_wrap(~Seasonality2,ncol = 5)+
  scale_shape_manual(values=c(21, 24))+
  theme_classic(base_size = 20)+
  theme(legend.position = "none")


#Figure SI-1
egg::ggarrange(EP_Seasonality_Spatial_struct_plot+rremove("x.title")+rremove("x.text"),TP_Seasonality_Spatial_struct_plot+rremove("x.title")+rremove("x.text"),TP_Seasonality_Spatial_struct_plot,
               ncol=1,nrow = 3)


EP_Seasonality_Spatial_struct_SD$Seasonality2<-as.factor(dplyr::recode(EP_Seasonality_Spatial_struct_SD$Seasonality, 
                                                                       "0.01" ="Seasonality = 0.01", 
                                                                       "0.2575" = "Seasonality = 0.2575",
                                                                       "0.505"= "Seasonality = 0.505",
                                                                       "0.7525"="Seasonality = 0.7525",
                                                                       "1"="Seasonality = 1"))


HS_Seasonality_Spatial_struct_SD$Seasonality2<-as.factor(dplyr::recode(HS_Seasonality_Spatial_struct_SD$Seasonality, 
                                                                       "0.01" ="Seasonality = 0.01", 
                                                                       "0.2575" = "Seasonality = 0.2575",
                                                                       "0.505"= "Seasonality = 0.505",
                                                                       "0.7525"="Seasonality = 0.7525",
                                                                       "1"="Seasonality = 1"))


TP_Seasonality_Spatial_struct_SD$Seasonality2<-as.factor(dplyr::recode(TP_Seasonality_Spatial_struct_SD$Seasonality, 
                                                                        "0.01" ="Seasonality = 0.01",
                                                                        "0.2575" = "Seasonality = 0.2575",
                                                                        "0.505"= "Seasonality = 0.505",
                                                                        "0.7525"="Seasonality = 0.7525",
                                                                        "1"="Seasonality = 1"))


EP_Seasonality_Spatial_struct_SD_plot<-ggplot(EP_Seasonality_Spatial_struct_SD,aes(x=Env_struct,y=Mean,group=Niche_diff))+
  #ylim(Min_Max_EP)+
  geom_errorbar(aes(ymin=LIC,ymax=UIC),width=.1)+
  geom_line(data = EP_Seasonality_Spatial_struct_SD,
            mapping = aes(x = Env_struct, y = Mean, group=Niche_diff))+
  ylab(expression('P.D. ( metacommunity-weighted mean'~italic(ep)~")"))+
  xlab("Seeded richness")+
  geom_point(aes(shape=Niche_diff),size=3,alpha=1,fill="white",stroke=2)+
  facet_wrap(~Seasonality2,ncol = 5)+
  scale_shape_manual(values=c(21, 24))+
  theme_classic(base_size = 20)+
  theme(legend.position = c(0.1, 0.9),
        legend.title=element_blank())


HS_Seasonality_Spatial_struct_SD_plot<-ggplot(HS_Seasonality_Spatial_struct_SD,aes(x=Env_struct,y=Mean,group=Niche_diff))+
  #ylim(Min_Max_EP)+
  geom_errorbar(aes(ymin=LIC,ymax=UIC),width=.1)+
  geom_line(data = HS_Seasonality_Spatial_struct_SD,
            mapping = aes(x = Env_struct, y = Mean, group=Niche_diff))+
  ylab(expression('P.D. ( metacommunity-weighted mean'~italic(hs)~")"))+
  xlab("Seeded richness")+
  geom_point(aes(shape=Niche_diff),size=3,alpha=1,fill="white",stroke=2)+
  facet_wrap(~Seasonality2,ncol = 5)+
  scale_shape_manual(values=c(21, 24))+
  theme_classic(base_size = 20)+
  theme(legend.position = "none")


TP_Seasonality_Spatial_struct_SD_plot<-ggplot(TP_Seasonality_Spatial_struct_SD,aes(x=Env_struct,y=Mean,group=Niche_diff))+
  #ylim(Min_Max_EP)+
  geom_errorbar(aes(ymin=LIC,ymax=UIC),width=.1)+
  geom_line(data = TP_Seasonality_Spatial_struct_SD,
            mapping = aes(x = Env_struct, y = Mean, group=Niche_diff))+
  ylab(expression('P.D. ( metacommunity-weighted mean'~italic(tp)~")"))+
  xlab("Seeded richness")+
  geom_point(aes(shape=Niche_diff),size=3,alpha=1,fill="white",stroke=2)+
  facet_wrap(~Seasonality2,ncol = 5)+
  scale_shape_manual(values=c(21, 24))+
  theme_classic(base_size = 20)+
  theme(legend.position = "none")


#Figure SI-II
egg::ggarrange(EP_Seasonality_Spatial_struct_SD_plot+rremove("x.title")+rremove("x.text"),HS_Seasonality_Spatial_struct_SD_plot+rremove("x.title")+rremove("x.text"),TP_Seasonality_Spatial_struct_SD_plot,
               ncol=1,nrow = 3)


EP_Competition<-partial_plots_rfscr_one_covariate3(formula=formula_fit_EP,Focal_Predictor="N_species",covariate="Comp_type",covariate2 ="Niche_diff",data=Data_simulations_final,repp=100)
HS_Competition<-partial_plots_rfscr_one_covariate3(formula=formula_fit_HS,Focal_Predictor="N_species",covariate="Comp_type",covariate2 ="Niche_diff",data=Data_simulations_final,repp=100)
TP_Competition<-partial_plots_rfscr_one_covariate3(formula=formula_fit_TP,Focal_Predictor="N_species",covariate="Comp_type",covariate2 ="Niche_diff",data=Data_simulations_final,repp=100)


EP_Competition_SD<-partial_plots_rfscr_one_covariate3(formula=formula_fit_EP_SD,Focal_Predictor="N_species",covariate="Comp_type",covariate2 ="Niche_diff",data=Data_simulations_final,repp=100)
HS_Competition_SD<-partial_plots_rfscr_one_covariate3(formula=formula_fit_HS_SD,Focal_Predictor="N_species",covariate="Comp_type",covariate2 ="Niche_diff",data=Data_simulations_final,repp=100)
TP_Competition_SD<-partial_plots_rfscr_one_covariate3(formula=formula_fit_TP_SD,Focal_Predictor="N_species",covariate="Comp_type",covariate2 ="Niche_diff",data=Data_simulations_final,repp=100)


EP_Competition_plot<-ggplot(EP_Competition,aes(x=N_species,y=Mean,group=Niche_diff))+
  geom_errorbar(aes(ymin=LIC,ymax=UIC),width=.1)+
  geom_line(data = EP_Competition,
            mapping = aes(x = N_species, y = Mean, group=Niche_diff))+
  ylab(expression('P.D. ( metacommunity-weighted mean'~italic(ep)~")"))+
  xlab("Seeded richness")+
  geom_point(aes(shape=Niche_diff),size=3,alpha=1,fill="white",stroke=2)+
  facet_wrap(~Comp_type,ncol = 5)+
  scale_shape_manual(values=c(21, 24))+
  theme_classic(base_size = 20)+
  theme(legend.position = c(0.1, 0.9),
        legend.title=element_blank())


HS_Competition_plot<-ggplot(HS_Competition,aes(x=N_species,y=Mean,group=Niche_diff))+
  geom_errorbar(aes(ymin=LIC,ymax=UIC),width=.1)+
  geom_line(data = HS_Competition,
            mapping = aes(x = N_species, y = Mean, group=Niche_diff))+
  ylab(expression('P.D. ( metacommunity-weighted mean'~italic(hs)~")"))+
  xlab("Seeded richness")+
  geom_point(aes(shape=Niche_diff),size=3,alpha=1,fill="white",stroke=2)+
  facet_wrap(~Comp_type,ncol = 5)+
  scale_shape_manual(values=c(21, 24))+
  theme_classic(base_size = 20)+
  theme(legend.position = "none")


TP_Competition_plot<-ggplot(TP_Competition,aes(x=N_species,y=Mean,group=Niche_diff))+
  geom_errorbar(aes(ymin=LIC,ymax=UIC),width=.1)+
  geom_line(data = TP_Competition,
            mapping = aes(x = N_species, y = Mean, group=Niche_diff))+
  ylab(expression('P.D. ( metacommunity-weighted mean'~italic(tp)~")"))+
  xlab("Seeded richness")+
  geom_point(aes(shape=Niche_diff),size=3,alpha=1,fill="white",stroke=2)+
  facet_wrap(~Comp_type,ncol = 5)+
  scale_shape_manual(values=c(21, 24))+
  theme_classic(base_size = 20)+
  theme(legend.position = "none")

#Figure SI-III
egg::ggarrange(EP_Competition_plot+rremove("x.title")+rremove("x.text"),TP_Competition_plot+rremove("x.title")+rremove("x.text"),TP_Competition_plot,
               ncol=1,nrow = 3)


EP_Competition_plot_SD<-ggplot(EP_Competition_SD,aes(x=N_species,y=Mean,group=Niche_diff))+
  #ylim(Min_Max_EP)+
  geom_errorbar(aes(ymin=LIC,ymax=UIC),width=.1)+
  geom_line(data = EP_Competition_SD,
            mapping = aes(x = N_species, y = Mean, group=Niche_diff))+
  ylab(expression('P.D. ( metacommunity-weighted mean'~italic(ep)~")"))+
  xlab("Seeded richness")+
  geom_point(aes(shape=Niche_diff),size=3,alpha=1,fill="white",stroke=2)+
  facet_wrap(~Comp_type,ncol = 5)+
  scale_shape_manual(values=c(21, 24))+
  theme_classic(base_size = 20)+
  theme(legend.position = c(0.1, 0.9),
        legend.title=element_blank())


HS_Competition_plot_SD<-ggplot(HS_Competition_SD,aes(x=N_species,y=Mean,group=Niche_diff))+
  #ylim(Min_Max_EP)+
  geom_errorbar(aes(ymin=LIC,ymax=UIC),width=.1)+
  geom_line(data = HS_Competition_SD,
            mapping = aes(x = N_species, y = Mean, group=Niche_diff))+
  ylab(expression('P.D. ( metacommunity-weighted mean'~italic(hs)~")"))+
  xlab("Seeded richness")+
  geom_point(aes(shape=Niche_diff),size=3,alpha=1,fill="white",stroke=2)+
  facet_wrap(~Comp_type,ncol = 5)+
  scale_shape_manual(values=c(21, 24))+
  theme_classic(base_size = 20)+
  theme(legend.position = "none")


TP_Competition_plot_SD<-ggplot(TP_Competition_SD,aes(x=N_species,y=Mean,group=Niche_diff))+
  #ylim(Min_Max_EP)+
  geom_errorbar(aes(ymin=LIC,ymax=UIC),width=.1)+
  geom_line(data = TP_Competition_SD,
            mapping = aes(x = N_species, y = Mean, group=Niche_diff))+
  ylab(expression('P.D. ( metacommunity-weighted mean'~italic(tp)~")"))+
  xlab("Seeded richness")+
  geom_point(aes(shape=Niche_diff),size=3,alpha=1,fill="white",stroke=2)+
  facet_wrap(~Comp_type,ncol = 5)+
  scale_shape_manual(values=c(21, 24))+
  theme_classic(base_size = 20)+
  theme(legend.position = "none")

#Figure SI-IV
egg::ggarrange(EP_Competition_plot_SD+rremove("x.title")+rremove("x.text"),HS_Competition_plot_SD+rremove("x.title")+rremove("x.text"),CON_Competition_plot_SD,
               ncol=1,nrow = 3)
```
