## Supplementary material for "Ecological selection of dispersal strategies in metacommunities: impact of landscape features and competitive dynamics": Code and Data to run simulations and generate figures: Functions and Simulations.R

```
library(ggplot2)
library(viridis)
library(WaveletComp)
library(vegan)
library(reshape2)
library(matrixStats)
library(ape)
library(ggpubr)
library(lattice)

#Simulation carried out in the manuscript: Ecological selection of dispersal strategies in metacommunities: impact of landscape features and competitive dynamics

####### Functions#####

#Emigration propensity function. It returns a vector showing species emigration propensity (from zero to 1) based on local performance
#Performance= Local performance considering habitat abiotic and biotic (competition) conditions.
#ep_responses = Species-specific "ep" parameter
#R.max= maximum intrinsic growth rate. It is equal across all species.


Emigration_propensity_function<-function(Perfomance,ep_responses,R.max){
  
  
  mu_emigration=ifelse(ep_responses>0,-1,1)
  
  #The lines below allow us to scale values of emigration propensity to range between 0 and 1 and the performance between 0 and 100.  
  Perfomance_scaled=(Perfomance*100)/R.max
  Min_response_emigration=mu_emigration*exp(ep_responses*100)
  Max_response_emigration=mu_emigration*exp(ep_responses*0)
  
  Focal_response_emigration=mu_emigration*exp(ep_responses*Perfomance_scaled)
  
  Scaled_response_Emigration_prob=1/(Max_response_emigration-Min_response_emigration)*(Focal_response_emigration-Max_response_emigration)+1
  
  
  return(Scaled_response_Emigration_prob)
  
}

#One can test this function and a produce similar figure as the one presented in the central panel
#of figure 1 using the code below

Testing_ep_levels<-seq(-0.1,0.1,length=10)

Figure_data<-sapply(1:length(Testing_ep_levels),function(x)
  Emigration_propensity_function(Perfomance = seq(0,3,length=1000),ep_responses = Testing_ep_levels[x],R.max = 3))


matplot(y=Figure_data,x=seq(0,3,length=1000),lty = 1,type = "l",xlab = "Performance",ylab = "Emigration Propensity")


#Habitat selection function. It returns a vector showing species attraction to a target patch (from zero to 1) based on target environmental suitability (which ranges from 0 to 1).
#Habitat_suit= Habitat suitabilit. (It only considers the match between species environmental optima and local environment).
#hs_responses = Species-specific "hs" parameter

Habitat_selection_function<-function(Habitat_suit,hs_responses){
  
  p_habitat_selection=ifelse(hs_responses<0,-1,1)
  
  #The lines below allow us to scale values of habitat selection to range between 0 and 1 
  
  
  Habitat_suit_scaled=(Habitat_suit*100)
  Max_response_immigration=p_habitat_selection*exp(hs_responses*100) 
  Min_response_immigration=p_habitat_selection*exp(hs_responses*0)
  
  Focal_response_immigration=p_habitat_selection*exp(hs_responses*Habitat_suit_scaled)
  
  
  Scaled_response_habitat_selection=1/(Max_response_immigration-Min_response_immigration)*(Focal_response_immigration-Max_response_immigration)+1
  
  
  return(Scaled_response_habitat_selection)
  
  
}

#One can test this function and a produce similar as the one presented in the central panel
#of figure 1 using the code below

Testing_hs_levels<-seq(-0.1,0.1,length=10)

Figure_data<-sapply(1:length(Testing_hs_levels),function(x)
  Habitat_selection_function(Habitat_suit = seq(0,1,length=1000),hs_responses = Testing_hs_levels[x]))


matplot(y=Figure_data,x=seq(0,1,length=1000),lty = 1,type = "l",xlab = "Performance",ylab = "Emigration Propensity")


# Traversal capacity function. It returns a N_site x N_site matrix showing species-specific traversal probability values between pairs of patches
#Coordinates_vl = Patch coordinates
#tp_responses = Species specific "tp" parameter

Traversal_capacity_function<-function(Coordinates_vl,tp_responses){
  
  
  focal_com<-as.matrix(exp(tp_responses*dist(Coordinates_vl)))
  
  #Scaling to range between 0 and 1
  Traversal_prob_min_dist<-exp(tp_responses*(60*sqrt(2)))# The maximum distance in a 60 by 60 landscape is given by the diagonal 60*sqrt(2)
  Traversal_prob_max_dist<-exp(tp_responses*0)
  
  Scaled_traversal_prob=1/(Traversal_prob_max_dist-Traversal_prob_min_dist)*(focal_com-Traversal_prob_max_dist)+1
  
  return(Scaled_traversal_prob)
  
}

#One can use the code below produce similar as the one presented in the central panel
#of figure 1 using the code below

Testing_tp_levels<-seq(-0.1,-0.99,length.out=10)


Figure_data<-sapply(1:length(Testing_tp_levels),function(x)
  exp(Testing_tp_levels[x]*c(0.0001,seq(1,60*sqrt(2),length=100))) )

matplot(y=Figure_data,lty = 1,type = "l",xlab = "Performance",ylab = "Emigration Propensity")


# Determines the match between species optimum and local env. conditions (ranges between 0 and 1)
#Env_cond = Local env conditions
#Sp_opt_conditions = Species optima given by parameter mu
#nbreadth = Niche breadth given by parameter sigma

habitat_suitability<-function(Env_cond,Sp_opt_conditions,nbreadth){
  exp(-((Env_cond - Sp_opt_conditions)^2/(2*nbreadth^2)))
}


# Densitiy-dependent competition.Returns a matrix giving the degree of  competition
#per species per site
#N = Population sizes
#alpha_intra= per-capta effect of a species on itself
#alpha_inter= per-capta effect of a species on others
competition<-function(N,alpha_intra,alpha_inter){
  competition_matrix<-matrix(0,nrow(N),ncol(N))
  for (c in 1:ncol(N)){
    for(d in 1:nrow(N)){
      Intra<-N[d,c]
      Inter<-abs(Intra-sum(N[d,]))
      competition_matrix[d,c]<-  1 / ( 1 + alpha_intra*Intra+alpha_inter*Inter)
    }
  }
  return(competition_matrix)
}

#Multivariate normal distribution to simmulate autocorrelated landscapes (See function Landscape_function)
rmvn <- function(n, mu = 0, V = matrix(1)) {
  p <- length(mu)
  if (any(is.na(match(dim(V), p)))) 
    stop("Dimension problem!")
  D <- chol(V)
  t(matrix(rnorm(n * p), ncol = p) %*% D + rep(mu, rep(n, p)))
}


#Function to simulate landscapes. It returns a list with two elements: 
#1-Coord: A matrix with patch xy coordinates; 
#2- Environment: A matrix N_sites x times showing local env conditions over time 

#N_sites= Number of sites, 
#time= Total time steps considered
#Env_sth: The level of environmental stochasticity.
#Seasonality= The magnitude (amplitude) of seasonal variations. The higher the value, ther higher the seasonality
#phi=  exponential decay of environmental similarity between pairs of patches as a function of their distances. The higher the value the more autocorrelated is the landscape

#If plot_landscapes = T, Generates a figure showing landscape at t1


Landscape_function<-function(N_sites,time,plot_landscape,Env_sth,phi,Seasonality){
  
  sites_names<-paste0("Site_",1:N_sites)
  
  
  for(ii in 1:10000){
    
    plot_coord=data.frame(Lat=runif(N_sites,0,60),Long=runif(N_sites,0,60))#Setting coordinates
    rownames(plot_coord)<-sites_names
    
    Elevation<-as.numeric(rmvn(1, rep(0,N_sites), exp(-phi * as.matrix(dist(plot_coord)))))
    w.small <- 1/as.matrix(dist(plot_coord))
    diag(w.small) <- 0
    Moran.I._sites<- Moran.I(Elevation, w.small)$observed
    
    if(Moran.I._sites>0){
      break
    }
    
    
    
    Attempt<-ii
    print(paste("Attempt #",Attempt, ":Generated Landscape is negatively autocorrelated. Trying again."))
  } 
  
  
  Elevation_grad<-(5-0)/(max(Elevation)-min(Elevation))*(Elevation-max(Elevation))+5#Rescaling values to 5
  
  
  
  Landscape<-matrix(0,nrow=N_sites,ncol=time)
  for(i in 1:length(Elevation)){
    Landscape[i,]=(Elevation_grad[i]+Seasonality*periodic.series(start.period = 12, length = time+3)+rnorm(time+3,0,Env_sth))[-1:-3]
  }
  
  Landscape_std<-as.matrix(5/(max(Landscape)-min(Landscape))*(Landscape-max(Landscape))+5) #Rescaling once again just in case
  row.names(Landscape_std)<-sites_names
  colnames(Landscape_std)<-paste0("T",seq (from= 1, to =time))
  
  
  
  #Generating a figure showing landscape at t1
  if(plot_landscape==T){
    
    Landscape_data<-data.frame(Lat=plot_coord$Lat,
                               Long=plot_coord$Long,
                               Env=Landscape_std[,1])
    
    
    Plot_Connect_step1<-ggplot(Landscape_data, aes(x = Lat, y = Long,fill=Env)) + 
      #geom_line(aes(group = gr),linetype="dashed", data = dat2)+
      ggtitle('Landscape in T1')+
      ylab("Y")+
      xlab("X")+
      #scale_size_continuous(range = c(1, 10))+
      geom_point(shape = 21, size = 5)+
      scale_fill_gradientn(colours = turbo(100),
                           limits=c(0,5))+
      #scale_size_continuous(range = c(0, 1))+
      coord_equal()+
      theme_bw()
    
    #print(Plot_Connect_step1)
    
    print(Plot_Connect_step1)
    
  }
  
  
  
  Results<-list(Landscape_std,plot_coord)
  names(Results)<-c("Environment","Coord")
  return(Results)
  
}


#Function to run metacommunity dynamics. It returns a lists with three elements
#1-A sites x species x time array representing the simulated metacommunity during the last seasonal cycle
#2-The metacommunitity-weighted mean (MWM) for ep, hs, tp
#3-The metacommunitity-weighted standard deviation (MWSD) for ep, hs, tp

#time= Total number of time steps
#N_sites= Total number of habitat patches in the landscape
#Env_struct= Level of spatial autocorrelation in habitat conditions. Values considered in this study are = seq(0.01,0.9,length=5)
#Seasonality= The magnitude (amplitude) of seasonal variations.  Values considered in simulations = seq(0.01,1,length=5)
#N_species=Number of species. Here we tested Species pools of sizes 50 and 300
#Nbreadth=Environmental tolerance (sigma)
#R.max=Maximum performance. Equal across all species
#Comp_types= Types of competitive dynamics considered. It can take the statues of "Stabilizing","Equalizing", and "Destabilizing"
#Niche_diff= A boolean stating Whether we assume  niche differentiation (TRUE) or not(FALSE), 

metacommunity_dynamics_and_dispersal_function<-function(
  time,
  N_sites,
  Env_struct,
  Seasonality,
  N_species,
  Nbreadth,
  R.max,
  Comp_types,
  Niche_diff
){
  
  ##Simulating Landscapes##
  sites_names<-paste0("Site_",1:N_sites)
  Landscape<-Landscape_function(N_sites = N_sites,time = time,phi=Env_struct,Env_sth = 0.1,plot_landscape = F,Seasonality = Seasonality)
  Landscape_env<-Landscape$Environment
  Coords<-Landscape$Coord
  
  ##Simulating species pools##
  sp_names<-paste0("Sp_",1:N_species)
  
  #Environmental optima according to Nich. Diff. assumption 
  if(Niche_diff==T){
  Species_opt<-seq(0,5,length=N_species)#Species optima under the assumption of Niche Differentiation
  }else{
  Species_opt<-rep(mean(Landscape$Environment),N_species)#Species optima under the assumption of neutrality
  }
  
  
  #Competition type tht will define the strength of co-specific and heterospecific competition
  if(Comp_types=="Stabilizing"){
    alphas=c(1/150,1/300)# Intra and inter alphas respectively 
  }else{
    if(Comp_types=="Equalizing"){
      alphas=c(1/200,1/200) 
    }else{
      alphas=c(1/300,1/150) 
    }
  }
  
  #Context depedent dispersal strategies
  ep_responses<-sample(seq(-0.1,0.1,length=N_species),replace = F)# Negative are species that stop emigrating as soon as performance improves a bit. Positive are those that only stop emigrating when conditions are very close to optimal
  hs_responses<-sample(seq(-0.1,0.1,length=N_species),replace = F)#Positive are species that are only attracted to optimal patches. Negative are the ones that go anywhere
  tp_responses<-sample(c(seq(-0.01,-0.99,length.out=N_species)),replace = F)# The more negative, the more species are constrained by the distance between sites
  
  # Generate array of N_site x N_site matrices x N_species dimensions. Each element of the array will be the traversal probability of a focal species to move
  #from a give site to other. This probability is given by species tp_responses and the euclidean distances between sites
  for(i in 1:dim(Connectivity_matrices)[3]){
    Connectivity_matrices[,,i]<-Traversal_capacity_function(Coordinates_vl =Coords,tp_responses = tp_responses[i] )
    Connectivity_matrices[,,i][Connectivity_matrices[,,i]<0]<-0# Fixing diagonals to 0 i.e. Probability ofcoming back to the natal patch just after emigrating from it = 0
  }
  
  
  # Generate array of N_site x N_Species matrices x time dimensions. Each element of the array will represent the metacommunity in a given moment in time.
  Communities<-array(0,dim=c(N_sites,N_species,time),
                     dimnames = list(c(rownames(Landscape$Environment)),c(sp_names),paste0("T",1:time)))
  
  t=1
  #Simulating metacommunity dynamics
  while(t <= time){
    
    print(paste("time",t))
    
    #habitat suitability for each species in each site at time t
    
    
    habitat_suitability_each_spxsite<-sapply(1:N_species,function(xxx)habitat_suitability(Landscape_env[,t],Species_opt[xxx],Nbreadth))
    colnames(habitat_suitability_each_spxsite)<-sp_names
    
    ### Seeding the model 
    if(t==1){
      Community_step1<-apply(Communities[,,t],2,function(x)rpois(length(x),0.5))
      row.names(Community_step1)<-row.names(Communities[,,t])
      
    }else{
      if(t%in%2:120){
        
        Seed<-do.call("cbind",lapply(1:N_species,function(z)rpois(N_sites,0.1)))
        Community_step1=Communities[,,t-1]+Seed
        row.names(Community_step1)<-row.names(Communities[,,t])
      }else{
        Community_step1=Communities[,,t-1]
        row.names(Community_step1)<-row.names(Communities[,,t])
      }
    }
    
  
    
    #Competition
    comp<-competition(Community_step1,alpha_intra = alphas[1],alpha_inter = alphas[2])
    
    
    
    
    #Calculating local performances based on habitat suitability and competition. 
    Performance_per_site_per_species<-R.max*comp*habitat_suitability_each_spxsite
    
    # Here Demographic stochasticity along with Performance 
    Community_step2<-matrix(0,nrow(Community_step1),ncol(Community_step1))
    dimnames(Community_step2)<-dimnames(Community_step1)
    for(l in 1:nrow(Community_step2)){
      Community_step2[l,]<-rpois(ncol(Community_step2),(Community_step1[l,]*Performance_per_site_per_species[l,]))
    }
    
    ##Dispersal###  
    
    #Emigration#
    
    #Defining emigration probability based on local performances 
    Emig_prob_per_species_per_site<-sapply(1:N_species,function(x)
      Emigration_propensity_function(Perfomance = Performance_per_site_per_species[,x],ep_responses = ep_responses[x],R.max = R.max))
    
    #Number of emigrants based on emigration probabilities (E_i_j_t)
    Emigrants_matrx<-sapply(1:N_species,function(x)rbinom(N_sites,Community_step2[,x],prob=Emig_prob_per_species_per_site[,x]))
    dimnames(Emigrants_matrx)<-dimnames(Community_step1)
    
    #Species leaving (emgirating) communities
    Community_step3<-Community_step2-Emigrants_matrx
    
    
    #Immigration#
    
    #The imigration matrix will store the number of immigrants from species i arriving at site j
    Immigration_matrx<-matrix(0,nrow(Community_step1),ncol(Community_step1))
    dimnames(Immigration_matrx)<-dimnames(Community_step1)
    #Creating a matrix showing the natal patch of dispersers ("Sites"), their ID ("Species"), and the number of dispersers ("Abund")
    Immigrants<-data.frame(which(Emigrants_matrx>0,arr.ind = T),Abund=Emigrants_matrx[which(Emigrants_matrx>0,arr.ind = T)])
    names(Immigrants)<-c("Sites","Species","Abund")
    
    # Defining patch attractiveness based on target habitat suitability
    Patch_attrac_per_species<-sapply(1:N_species,function(x)
      Habitat_selection_function(Habitat_suit = habitat_suitability_each_spxsite[,x],hs_responses = hs_responses[x]))
    
    #Immigration probability (Disp_prob) is given species-specific patch attractiveness * species specific traversal probability.
    #The Disp_prob is a Sites X Sites X Species array, showing the probability of species to arrive in a given site based on its habitat quality (defined by their habitat selection trait) and distance from focal site (defined by their traversal capacity trait).
    Disp_prob<-sapply(1:N_species,function(x)Connectivity_matrices[,,x]*Patch_attrac_per_species[,x],simplify = "array")
    
    
    for(z in 1:nrow(Immigrants)){
      Destination<-table(sample(sites_names[-Immigrants[z,"Sites"]],size=Immigrants[z,"Abund"],replace = T,prob=Disp_prob[-Immigrants[z,"Sites"],Immigrants[z,"Sites"],Immigrants[z,"Species"]]))
      
      Immigration_matrx[names(Destination),Immigrants[z,"Species"]]<-Immigration_matrx[names(Destination),Immigrants[z,"Species"]]+as.numeric(Destination)
    }
    
    Community_step4<-Community_step3+Immigration_matrx
    
    
    Communities[,,t]<-Community_step4
    
    t=t+1
    
  }#End of simulations
  
  MC_last_Seasonal_Cycle<-Communities[,,c((time-11):time)]
  
  Final_metacom<-rowMeans(MC_last_Seasonal_Cycle, dims = 2)
  Final_metacom_occupancy<-decostand(Final_metacom,"pa")
  
  Peristant_species<-which(colSums(Final_metacom)>0)
  
  #Dominance is estimated based on species regional abundances and occupancy
  Dominance<-colSums(Final_metacom[,Peristant_species])*(colSums(Final_metacom_occupancy[,Peristant_species])/N_sites)
  
  Mean_tp<-weighted.mean(tp_responses[Peristant_species],w = Dominance)
  Mean_ep<-weighted.mean(ep_responses[Peristant_species],w = Dominance)
  Mean_hs<-weighted.mean(hs_responses[Peristant_species],w = Dominance)
  
  
  SD_tp<-weightedSd(tp_responses[Peristant_species],w = Dominance)
  SD_ep<- weightedSd(ep_responses[Peristant_species],w = Dominance)
  SD_hs<-weightedSd(hs_responses[Peristant_species],w = Dominance)
  
  Dominant_traits<-c(Mean_ep,Mean_hs,Mean_tp)
  names(Dominant_traits)<-c("MWM-ep","MWM-hs","MWM-tp")
  
  Trait_diversity<-c(SD_ep,SD_hs,SD_tp)
  names(Trait_diversity)<-c("MWSD-ep","MWSD-hs","MWSD-tp")
  
  Results<-list(MC_last_Seasonal_Cycle,
                Dominant_traits,
                Trait_diversity)
  
  return(Results)
  
}


######Example (it can take a few minutes to run)######

##Landscape parameters#
time<-1200# Total number of time steps
N_sites<-50
Env_struct<-c(0.9)#Phi Values considered in simulations = seq(0.01,0.9,length=5)
Seasonality<-0.1# Values considered in simulations = seq(0.01,1,length=5)


#Creating Illustrative landscape
Landscape<-Landscape_function(N_sites = N_sites,time = time,phi=Env_struct,Env_sth = 0.1,plot_landscape = T,Seasonality = Seasonality)
Landscape_env<-Landscape$Environment
matplot(t(Landscape_env[,1:100]),type = "l")#Variation in Env. over time


##Seeded Species pools parameters###
N_species<-50# Number of species. Here we tested Species pools of sizes 50 and 300
Nbreadth<-1# Environmental tolerance
R.max<-3#Maximum performance. Equal across all species
Comp_types<-"Stabilzing"# Type of competition (Stabilizing,"Equalizing" and "Destabilizing") 

Simulation_results<-metacommunity_dynamics_and_dispersal_function(time=time,
                                              N_sites=N_sites,
                                              Env_struct=Env_struct,
                                              Seasonality=Seasonality,
                                              N_species=N_species,
                                              Nbreadth=Nbreadth,
                                              R.max=R.max,
                                              Comp_types=Comp_types,
                                              Niche_diff = T)
```
