## Supplementary material for "Ecological selection of dispersal strategies in metacommunities: impact of landscape features and competitive dynamics": Code and Data to run simulations and generate figures: README.docx

Title: The ecological selection of metacommunity dynamics on context-dependent dispersal strategies: The role of landscape features and competition dynamics

Brief Description

While the influence of dispersal on metacommunities is subject of intense research, we still do not understand how species-species and species-environment relationships determine the success of different dispersal strategies in metacommunities. To address this, we employed simulation models considering species with distinct context-dependent dispersal strategies involved in the three stages of dispersal (departure, transience, and settlement). These species were allowed to reach coexistence at the metacommunity scale under various competitive hierarchies and different levels of spatial and temporal environmental variability. By assessing the context-dependent dispersal strategies of species that persisted and dominated metacommunities, we could understand how metacommunity dynamics impose ecological selection on dispersal.

This folder contains:

1- Functions and Simulations Notated.R- Code to run simulation models, assess metacommunity-weighted mean and standard deviation of species dispersal traits (*ep, hs*, and *tp*).
2- Analyses and Figures.R- Code to run Random Forest models, generate heatmaps, and partial plots. All figures in the manuscript and SI can be generated using the code in this manuscript.
3- Data_Simulations_Raw.txt. A 6000 x 11 data file with the results of simulations performed using code in provided in Functions and Simulations Notated.R. Each line contains one of the 20 replicates X 5 levels of seasonality (values described in line 459 of Functions and Simulations Notated.R) X 5 levels of spatial structure of environmental conditions (values described in line 458 of Functions and Simulations Notated.R) X 2 species pool sizes (50 and 300 species) X 3 competition structures (stabilizing, equalizing, destabilizing) X 2 Niche differentiation assumptions (Niche Diff, Neutrality).
4- Data_Simulations_Final.txt. A 300 x 11 data file with the average results of the cross factorial combination of simulation scenarios across the 200 replicates. All the analyses and figures reported in this study were generated using this dataset.

Data files description

Data_Simulations_Raw.txt.

A 6000 x 11 data file with the results of simulations performed using code in provided in Functions and Simulations Notated.R. Each line contains one of the 20 replicates X 5 levels of seasonality (values described in line 459 of Functions and Simulations Notated.R) X 5 levels of spatial structure of environmental conditions (values described in line 458 of Functions and Simulations Notated.R) X 2 species pool sizes (50 and 300 species) X 3 competition structures (stabilizing, equalizing, destabilizing) X 2 Niche differentiation assumptions (Niche Diff, Neutrality).

Each row is the results of a replicate of each simulation scenario. This raw data is provided to aid with the reproducibility of the results reported in this manuscript.

Column IDs and description:

N_species = Number of species seeded in the landscape at t1

Env_struct= Level of spatial autocorrelation in environmental conditions

Seasonality= Degree of seasonality in the landscape

Comp_type = Type of competitive structure

Niche_diff = Assumption regarding species ecological equivalence

Avg.Weighted_EP = Metacommunity_Weighted_mean ep

Avg.Weighted_HS = Metacommunity_Weighted_mean hs

Avg.Weighted_TP = Metacommunity_Weighted_ mean tp

SD_EP = Metacommunity_Weighted_standard deviation ep

SD_HS = Metacommunity_Weighted_standard deviation hs

SD_TP = Metacommunity_Weighted_standard deviation TP

Data_Simulations_Final.txt.

A 300 x 11 data file with the average results of the cross factorial combination of simulation scenarios across the 200 replicates. All the analyses and figures reported in this study were generated using this dataset.

Each row represents the average value obtained across the 20 replicates of a simulation scenario.

This is the data file used to generate the figures and run the analyses reported in the main manuscript.

Column IDs and description:

N_species = Number of species seeded in the landscape at t1

Env_struct= Level of spatial autocorrelation in environmental conditions

Seasonality= Degree of seasonality in the landscape

Comp_type = Type of competitive structure

Niche_diff = Assumption regarding species ecological equivalence

Avg.Weighted_EP = Metacommunity_Weighted_mean ep (average across 20 replicates)

Avg.Weighted_HS = Metacommunity_Weighted_mean hs (average across 20 replicates)

Avg.Weighted_TP = Metacommunity_Weighted_ mean tp (average across 20 replicates)

SD_EP = Metacommunity_Weighted_standard deviation ep (average across 20 replicates)

SD_HS = Metacommunity_Weighted_standard deviation hs (average across 20 replicates)

SD_TP = Metacommunity_Weighted_standard deviation TP (average across 20 replicates)
